# Statistical mining of triple-negative breast cancer-specific nanobodies among huge libraries from immunized alpacas

**DOI:** 10.1101/2023.02.23.529685

**Authors:** Ryota Maeda, Hiroyuki Yamazaki, Ryoga Kobayashi, Seishiro Yamamoto, Kazuki Kasai, Akihiro Imura

## Abstract

Breast cancer can be classified into several types according to the expression patterns of human epidermal growth factor receptor 2 (Her2), oestrogen receptor (ER), and progesterone receptor (PgR) proteins. The prognosis of patients with tumors showing low Her2 expression and no ER and PgR expression—categorized as triple-negative breast cancer (TNBC)—is worst among these groups. Due to the lack of specific antibodies for TNBC, curative treatments for TNBC remain limited. Antibodies targeting TNBC have potential as diagnostic and therapeutic tools. Here, we generate a panel of nanobodies targeting TNBC cell lines by immunizing alpacas and subsequently panning the resulting phage libraries with TNBC cell lines. We show that several clones exclusively stain Her2-negative cells in tissues of breast cancer patients, and a few clones stain both Her2-positive and Her2-negative regions in these tissues. These clones can be applied to patient-specific therapies using drug-conjugated antibodies, radiolabelled antibodies, chimaera antigen receptor T cells, or drug delivery components, as well as to TNBC diagnosis.

## Introduction

Breast cancer is the most common cancer in women. In 2022, more than 0.4 million women had breast cancer, and at least 43,000 breast cancer patients died in the US^1^. Clinically, breast cancer is a wide variety of diseases^2–6^; treatments are targeted towards molecular markers, namely, oestrogen receptor (ER), progesterone receptor (PgR), and human epithelial growth factor receptor 2 (Her2). However, 10-20% of all breast cancers with no expression of the ER and PgR and lacking amplification of Her2 are categorized as triple-negative breast cancer (TNBC)^7^. TNBC exhibits the most aggressive course of the breast cancers, and TNBC patients have a higher rate of distant recurrence and a poorer prognosis^8^. TNBC occurs mostly in premenopausal women under the age of 40, representing 25-30% of breast cancers in this age group^9^. Endocrine therapy was effective for breast cancer patients with tumours that express hormone receptors (ER or PgR). Targeted therapy using anti-Her2 antibodies benefited patients with aberrantly Her2-expressing cancer^10–12^. However, due to the lack of target molecules, patients with TNBC are not sensitive to endocrine therapy or molecular targeted therapy^13^. Chemotherapy is the first-choice treatment, and clinical studies of immunotherapy for inhibiting programmed cell death-1 (PD-1) are ongoing; however, their efficacy and the overall survival rate of TNBC are still poor^14, 15^. In addition, it is thought that tumour recurrences originate from residual metastatic lesions. Therefore, it is urgent to develop new targets for TNBC to fulfil these unmet medical needs.

To establish monoclonal anticancer antibodies, it has been widely attempted to immunize animals with cancer tissues or cancer-derived cell lines^16–22^. Since camelids, including alpacas, have heavy chain-only antibodies—single variable domain on a heavy chain (VHH), or nanobodies—they are well suited to phage-display biopanning, and the genes encoding the nanobodies are easily sequenced^23–28^. We used the TNBC cell lines to immunize alpacas, performed bio-panning using the TNBC cells, and identified TNBC cell line-specific nanobodies. Using microscopic analysis with a panel of breast cancer tissues, we show that these nanobodies targeted breast cancer that was not highly expressing Her2. We used TNBC cell lines to immunize alpacas, performed biopanning using TNBC cells, and identified TNBC cell line-specific nanobodies. Using immunostaining analysis of a panel of breast cancer tissues, we show that these nanobodies targeted breast cancers that did not express ER, PgR, and high levels of Her2.

## Results

### Breast cancer cell lines used for immunizing alpacas

We immunized alpacas with the well-characterized breast cancer cell lines: SK-BR-3, MDA-MB-231, T47D, BT-474, HS578T, BT-549, MDA-MB-436, MDA-MB-468, and HCC1937^29, 30^. Her2 is highly expressed in SK-BR-3 and BT-474 cells, and ER and PgR are expressed in T47D and BT-474 cells^31, 32^. Other cell lines—MDA-MB-231, HS578T, BT-549, MDA-MB-436, MDA-MB-468, and HCC1937—express neither Her2 nor the hormone receptors and are considered TNBC cell lines. Subconfluent cultured cells were gently detached from culture dishes, washed, and fixed with paraformaldehyde (PFA). We immunized an adult alpaca five times at 2-week intervals and collected peripheral blood cells after the fourth and fifth injections.

### Cell-based biopanning

We generated phage libraries from the VHH genes after the fourth injection and after the fifth injection. The two resulting mother libraries were precleared by incubation with nonbreast cancer cell lines, including human embryonic kidney (HEK) 293 cells, human carcinoma of cervix-derived HeLa cells, African green monkey-derived COS-7 cells, human epidermoid carcinoma-derived A-431 cells, and human hepatoblastoma-derived HepG2 cells. Second, the precleared mother libraries were panned with the breast cancer cell lines SK-BR-3, MDA-MB-231, HS578T, BT-549, MDA-MB-436, MDA-MB-468, HCC1937, and HCC1599, among which HCC1599 cells are TNBC cells and had not been used to immunize the alpacas. Third, we read the VHH sequences in each biopanned sublibrary using a next-generation sequencer. Finally, we selected clones that were enriched by the TNBC cell lines (MDA-MB-231, HS578T, BT-549, MDA-MB-436, MDA-MB-468, HCC1937, and HCC1599); we then expressed 17 clones in Escherichia coli for further analysis.

We found that several VHH clones, TNBC21, TNBC72, TNBC172, TNBC242, and TNBC259, were enriched only in the MDA-MB-231 cell-panned sublibraries and that TNBC673, TNBC769, and TNBC3511 were enriched only by MDA-MB-468; TNBC610 and TNBC13289 were slightly enriched by HCC1937 (Figure 1a). Another 7 clones were enriched among HCC1599 cell-panned sublibraries as well (i.e., sublibraries generated with the cell line that had not been used for immunization).

**Figure 1.**
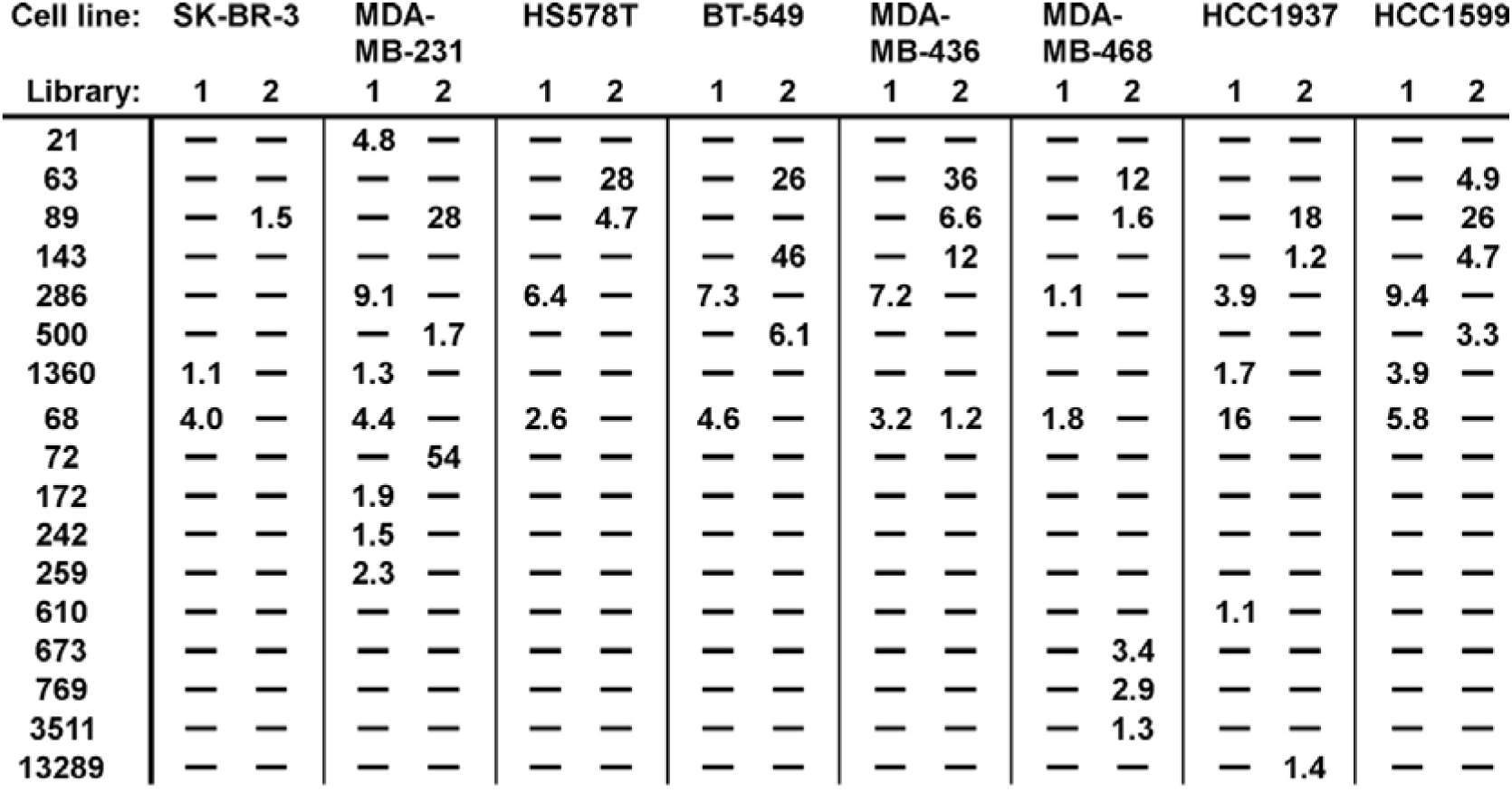
Clone enrichment scores. Comparison of enrichment scores after panning with the indicated cell lines. Inversed *P* values on the log scale are shown. SK-BR-3 cells are Her2 positive; other cells are triple-negative breast cancer cells. HCC1599 cells were not used for immunization. -), not enriched.

TNBC68 was enriched in all cell-panned sublibraries; TNBC89 was enriched in all sublibraries except for the BT-549 cell-panned sublibrary. TNBC286 was enriched in all TNBC cell-panned sublibraries and not in the Her2-expressing SK-BR-3 cell-panned sublibraries. The 17 selected clones had complementary determinant regions (CDRs) of different lengths and sequences (Figure 1b). We further assessed TNBC21, TNBC63, TNBC89, TNBC143, TNBC286, TNBC500, and TNBC1360 clones in a single monomer and the other 10 clones in a linked tandem dimer.

### Staining breast cancer cell lines

For immunostaining analysis, we prepared SK-BR-3 cells as the Her2-positive cell line; we used MDA-MB-231, HS578T, BT-549, and MDA-MB-436 cells as the mesenchymal-like TNBC cell lines; and we used MDA-MB-468, HCC1937, and HCC1599 cells as the basal-like TNBC cell lines. We scored the staining images into four levels: strongly positive; moderately positive; weakly positive, partially (<10%) positive, or only dividing cells positive; and negative (Figure 2 and Supplementary Figures 1 and 2). TNBC143 stained five TNBC cell lines but not Her2-positive SK-BR-3 cells. TNBC89, TNBC72, and TNBC769 stained mesenchymal-like and basal-like TNBC cell lines. TNBC63 and TNBC172 stained mesenchymal-like TNBC cell lines; in contrast, TNBC242, TNBC259, TNBC610, TNBC673, TNBC1360, TNBC3511, and TNBC13289 stained basal-like TNBC cell lines. Although TNBC21, TNBC68, and TNBC500 appeared to be enriched upon biopanning, they did not stain any cells tested well, probably due to high background signal.

**Figure 2.**
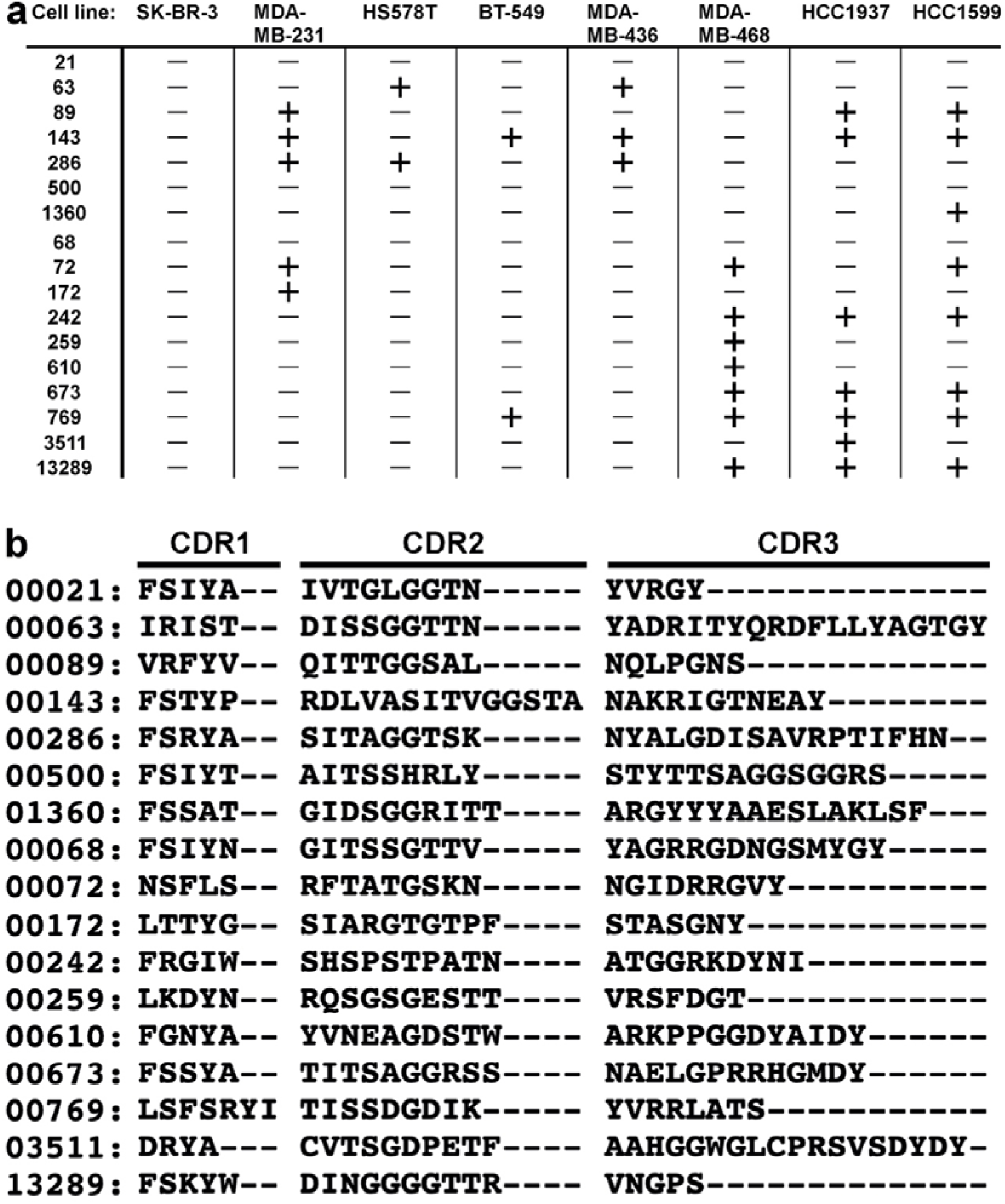
Staining of the cell lines and sequence alignment. **a**, Staining results for each clone in the cell lines are shown. -), negative or weak signals, and +), positive or strong signals. The corresponding raw data are available in Supplementary Figure 1. **b**, Alignments of the CDR1, CDR2, and CDR3 sequences of the selected 17 clones.

### Staining breast cancer tissues

Having tested immunostaining in cultured cells, we next applied the clones to immunohistochemical staining (red) of frozen breast cancer tissue sections (Frozen Tissue array—37 different breast tumours and three corresponding normal controls: T6235086-5, C105028, BioChain). We counterstained for Her2 using an anti-Her2 antibody (green) and nucleic acid with DAPI (blue); we then imaged the sections under the same conditions using confocal laser microscopy. In invasive ductal carcinoma tissues, we found that TNBC143 stained the tissues in four patterns: i) Her2-negative basal cells surrounding Her2-positive apical cells (A1-1 and D2-1 of Figure 3a), ii) ductal cells with high levels of chromatin (A3-2 of Figure 3a and Supplementary Figure 3), iii) marginal Her2-negative cells neighbouring Her2-amplified cells (B8-2 and C8-2 of Figure 3a), and iv) Her2-negative stromal cells (C3-2 and C5-1 of Figure 3a). No or weak signals for TNBC143 were observed in normal breast tissues (Figure 3b). We scored the staining results according to the pathological criteria: ratio of positive cells (0; none, 1; less than 1%, 2; between 1 and 10%, 3; between 10 and 50%, and 4; more than 50%) and intensity (0; negative, 1; weak, 2; moderate, and 3; strong). When the sum of both scores above 3 was defined as positive staining, 28 out of 37 breast cancer samples were TNBC143-positive.

**Figure 3.**
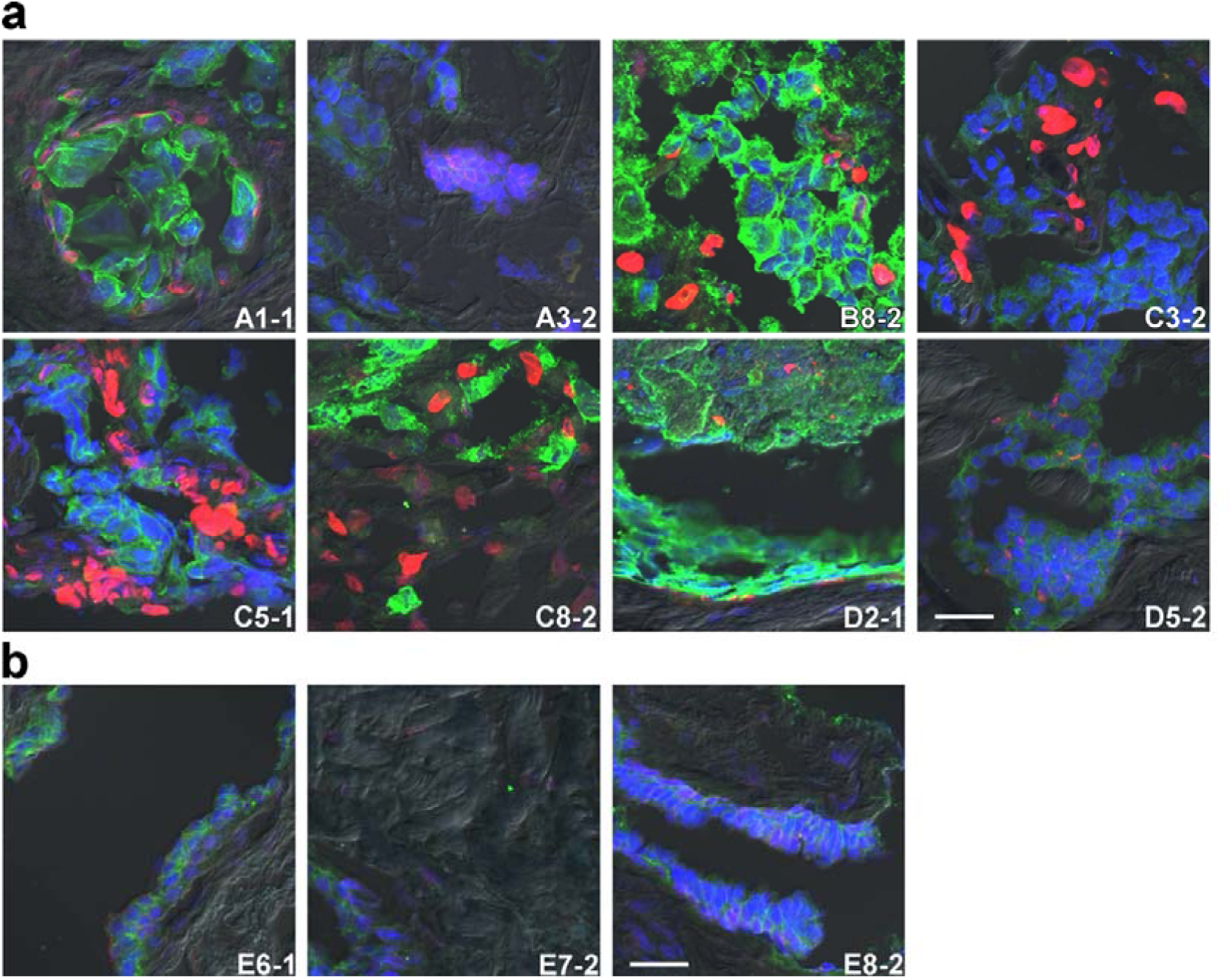
Staining patterns of breast cancer tissues with the TNBC143m clone. **a**, Representative images of tissue staining were merged; clones are indicated in red. Her2, green colour. **b**, Staining of the healthy breast tissue samples is shown. Sample IDs and observed field numbers are inset. Scale bars = 26.2 μm. Unmerged images of A3-2 are shown in Supplementary Figure 3.

Similar to TNBC143, TNBC89 stained invasive ductal carcinoma tissues in four patterns: i) basal cells surrounding Her2-positive apical cells (A1-2 and B2-2 of Figure 4a), ii) aggressive ductal cells (A3-2 and B7-3 of Figure 4a and Extended Data Figure 3), iii) marginal cells (B5-1 and C2-1 of Figure 4a), and iv) stromal cells (A6-2 and C5-1 of Figure 4a). TNBC89 weakly stained basal cells of the duct in normal breast tissues (Figure 4b). TNBC673 yielded similar staining patterns: i) basal cells (E2-1), ii) aggressive ductal cells (B7-1 and D6-1), iii) marginal cells (A7-2 and B4-1), and iv) stromal cells (C3-2, C5-1, and C6-1) (Figure 5a and Extended data Figure 3); TNBC673 weakly stained basal cells of the duct in the normal breast tissues (Figure 5b).

**Figure 4.**
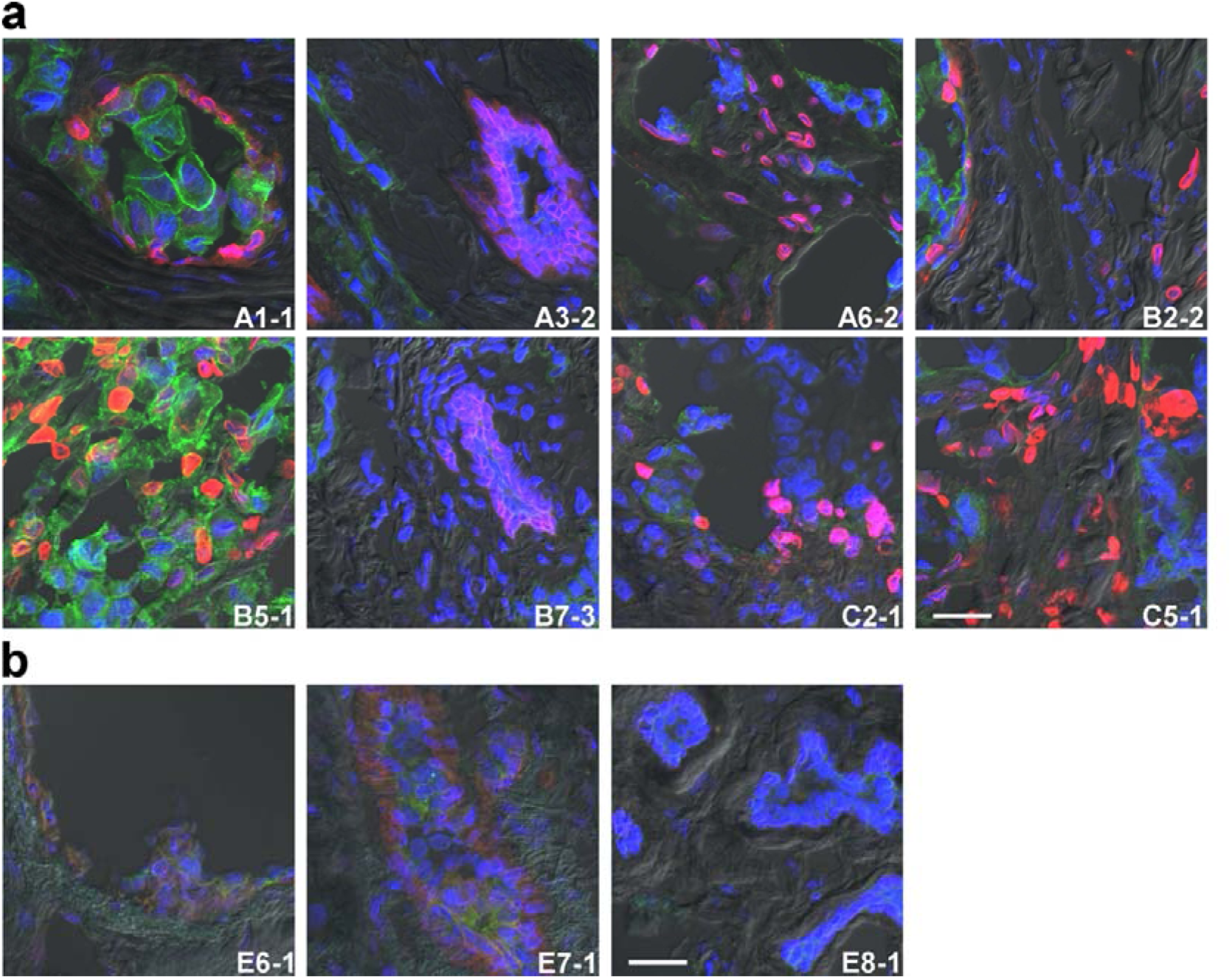
Staining patterns of breast cancer tissues with the TNBC89m clone. **a**, Representative images of tissue staining were merged; clones are indicated in red. Her2, green colour. **b**, Staining of healthy breast tissue samples is shown. Sample IDs and observed field numbers are inset. Scale bars = 26.2 μm. Unmerged images of A3-2 and B7-3 are shown in Supplementary Figure 3.

**Figure 5.**
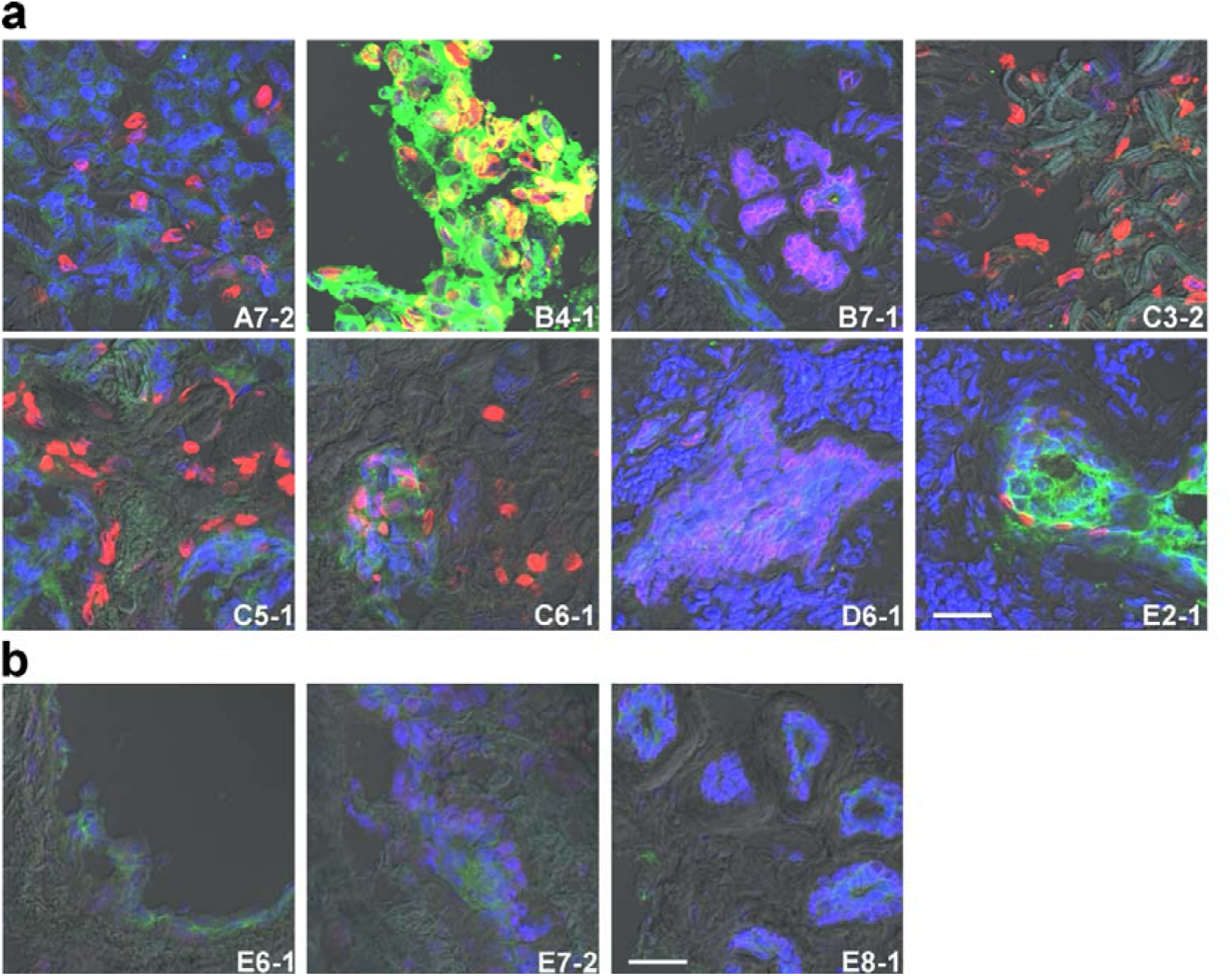
Staining patterns of breast cancer tissues with the TNBC673m clone. **a**, Representative images of tissue staining were merged; clones are indicated in red. Her2, green colour. **b**, Staining of healthy breast tissue samples is shown. Sample IDs and observed field numbers are inset. Scale bars = 26.2 μm. Unmerged images of D6-1 are shown in Supplementary Figure 3.

Although only weak signals were observed in the cell lines tested, TNBC68 significantly stained breast cancer tissues (Figure 6). In contrast to the above clones, TNBC68 stained a portion of the basal cells in invasive ductal carcinomas (C1-2 and D4-1 of Figure 6a and Extended Data Figure 4). TNBC68 stained marginal cells (A4-2, D8-1, and E1-1 of Figure 6a and Extended Data Figure 4) and stromal cells (A2-2 and E4-1 of Figure 6a and Extended Data Figure 4). TNBC68 also stained basal cells of the milk duct in normal breast tissues (Figure 6b).

**Figure 6.**
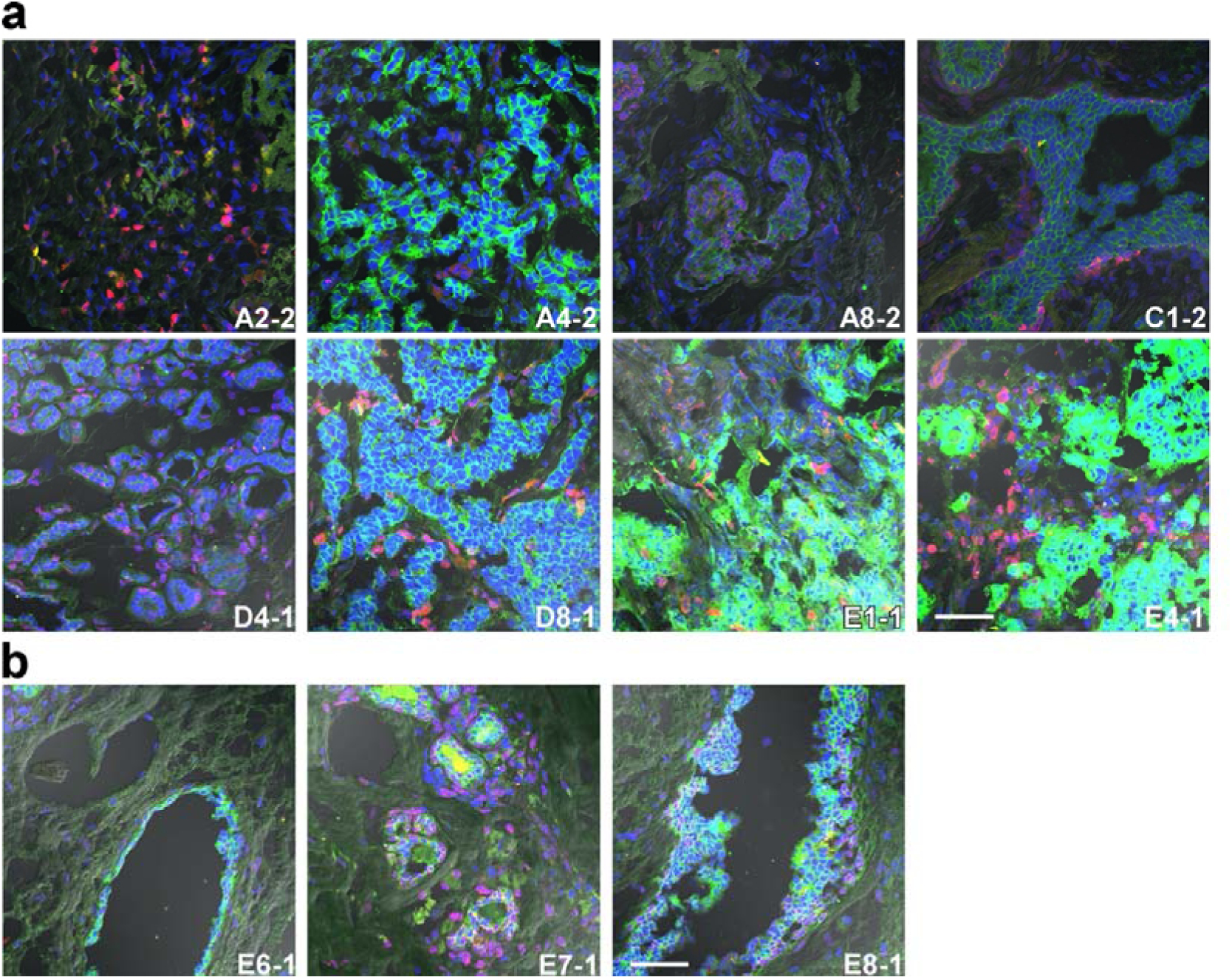
Staining patterns of breast cancer tissues with the TNBC68d clone. **a**, Representative images of tissue staining were merged; clones are indicated in red. Her2, green colour. **b**, Staining of healthy breast tissue samples is shown. Sample IDs and observed field numbers are inset. Scale bars = 63.7 μm. Unmerged images of A4-2, A8-2, C1-2, D4-1, and E1-1 are shown in Supplementary Figure 4.

TNBC242 stained mesenchymal TNBC cell lines and most intensively stained invasive ductal carcinoma tissues (Figure 7 and Extended Data Figure 5). In normal breast cancer tissues, TNBC242 stained Her2-positive apical cells of the mammary gland (Figure 7b). In invasive ductal carcinoma tissues, TNBC242 stained Her2-negative ducts, including basal cells (C3-1, D2-1, and D3-2) and aggressive ductal cells (D1-1 and D4-1). Most impressively, TNBC242 strongly stained stromal regions with no expression of Her2 and no tubular structures (C2-2, C5-2, and C8-2).

**Figure 7.**
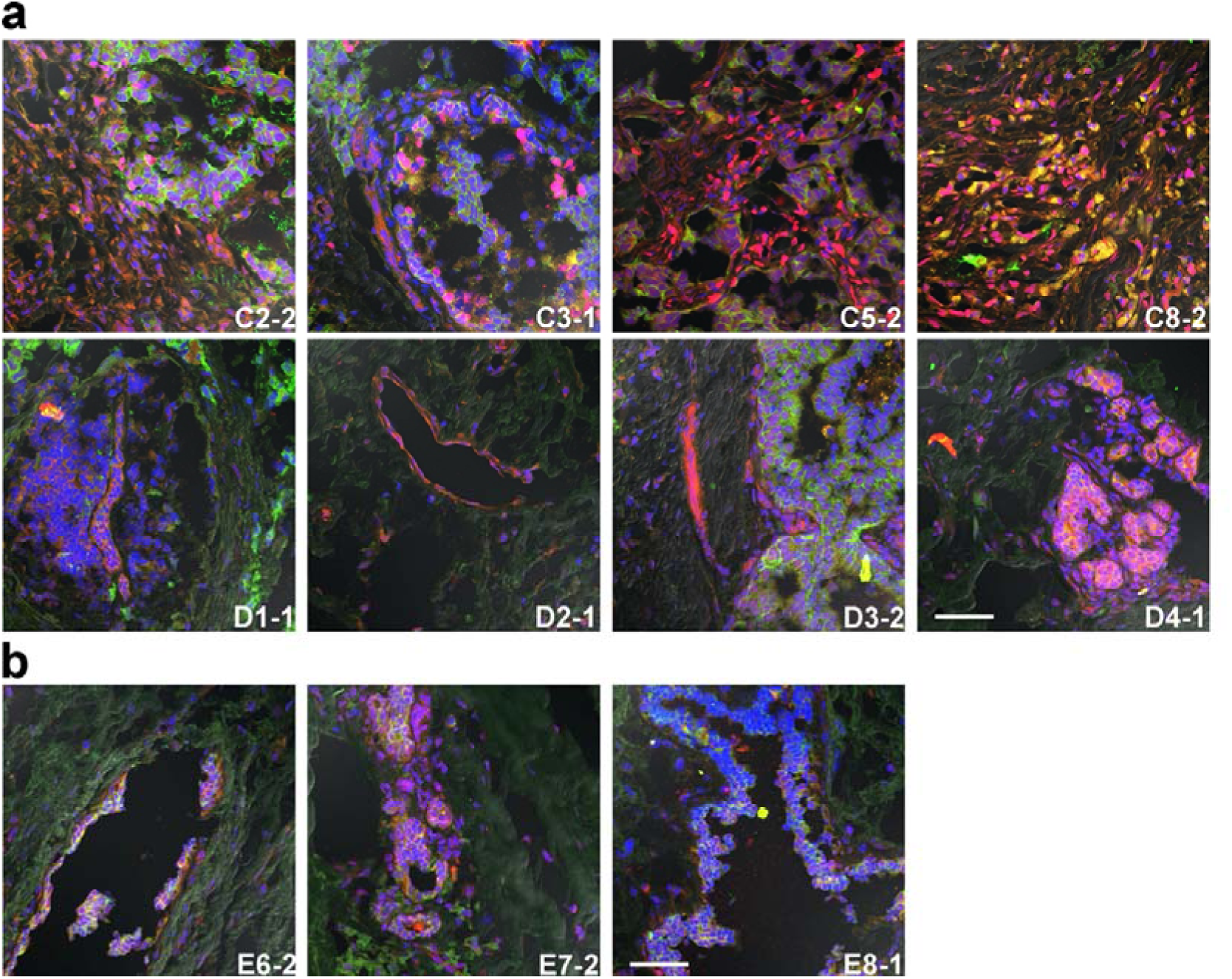
Staining patterns of breast cancer tissues with the TNBC242d clone. **a**, Representative images of tissue staining were merged; clones are indicated in red. Her2, green colour. **b**, Staining of healthy breast tissue samples is shown. Sample IDs and observed field numbers are inset. Scale bars = 63.7 μm. Unmerged images of C2-2, C3-1, D1-1, D2-1, D3-2, and D4-1 are shown in Supplementary Figure 5.

TNBC610 stained only one mesenchymal TNBC cell line, MDA-MB-468, and stained breast cancer tissues in three patterns: a few cells neighbouring Her2-positive cells (A6-1, B6-1, C7-1, and C8-1 of Figure 8a), basal cells of aggressive Her2-positive ductal cells (B7-2 and E5-2 of Figure 8a and Extended data Figure 6), and stromal cells of Her2-negative regions (C5-1 and C8-1 of Figure 8a). No TNBC610 signals were observed in normal breast tissues (Figure 8b). Finally, TNBC13289, which stained only mesenchymal TNBC cell lines, gave mainly sporadic signals neighbouring Her2-positive ductal cells in invasive ductal carcinoma tissues (B2-2, C3-2, D4-2, C5-2, E2-1, and E3-1 of Figure 9a and Extended Data Figure 6). In a few cases, TNBC13289 stained Her2-amplified apical cells (E1-1). TNBC13289 weakly stained Her2-positive apical cells of the mammary glands in normal breast tissue (Figure 9b). Finally, we counterstained ER and PgR using anti-ER and anti-PgR antibodies (green). We found that TNBC89 stained invasive ductal carcinoma tissues where both ER and PgR were negative (Figure 10 and Extended Data Figure 7).

**Figure 8.**
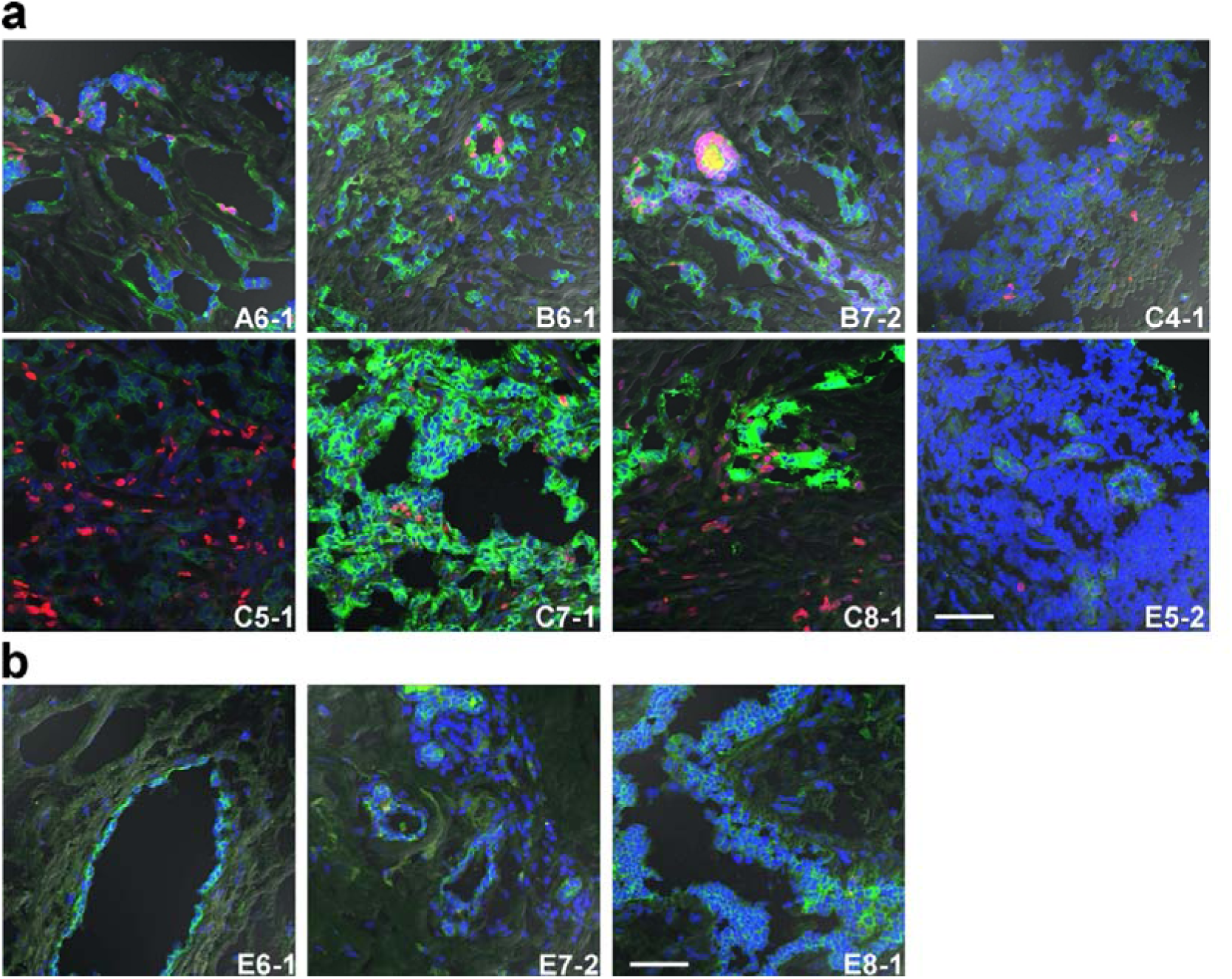
Staining patterns of breast cancer tissues with the TNBC610d clone. **a**, Representative images of tissue staining were merged; clones are indicated in red. Her2, green colour. **b**, Staining of healthy breast tissue samples is shown. Sample IDs and observed field numbers are inset. Scale bars = 63.7 μm. Unmerged images of B7-2, C7-1, and E5-2 are shown in Supplementary Figure 6.

**Figure 9.**
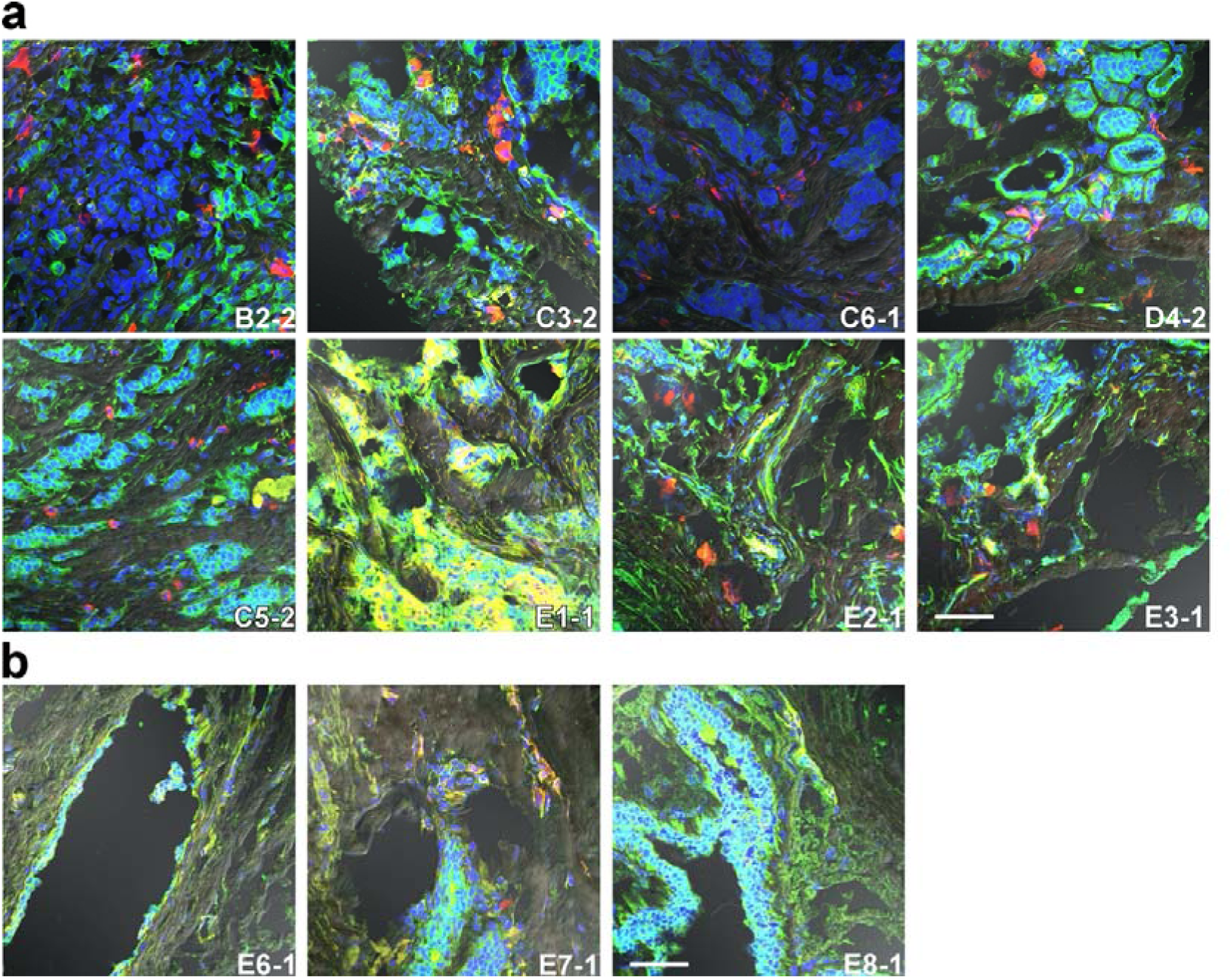
Staining patterns of breast cancer tissues with the TNBC13289d clone. **a**, Representative images of tissue staining were merged; clones are indicated in red. Her2, green colour. **b**, Staining of healthy breast tissue samples is shown. Sample IDs and observed field numbers are inset. Scale bars = 63.7 μm. Unmerged images of C3-2, E1-1, and E3-1are shown in Supplementary Figure 6.

**Figure 10.**
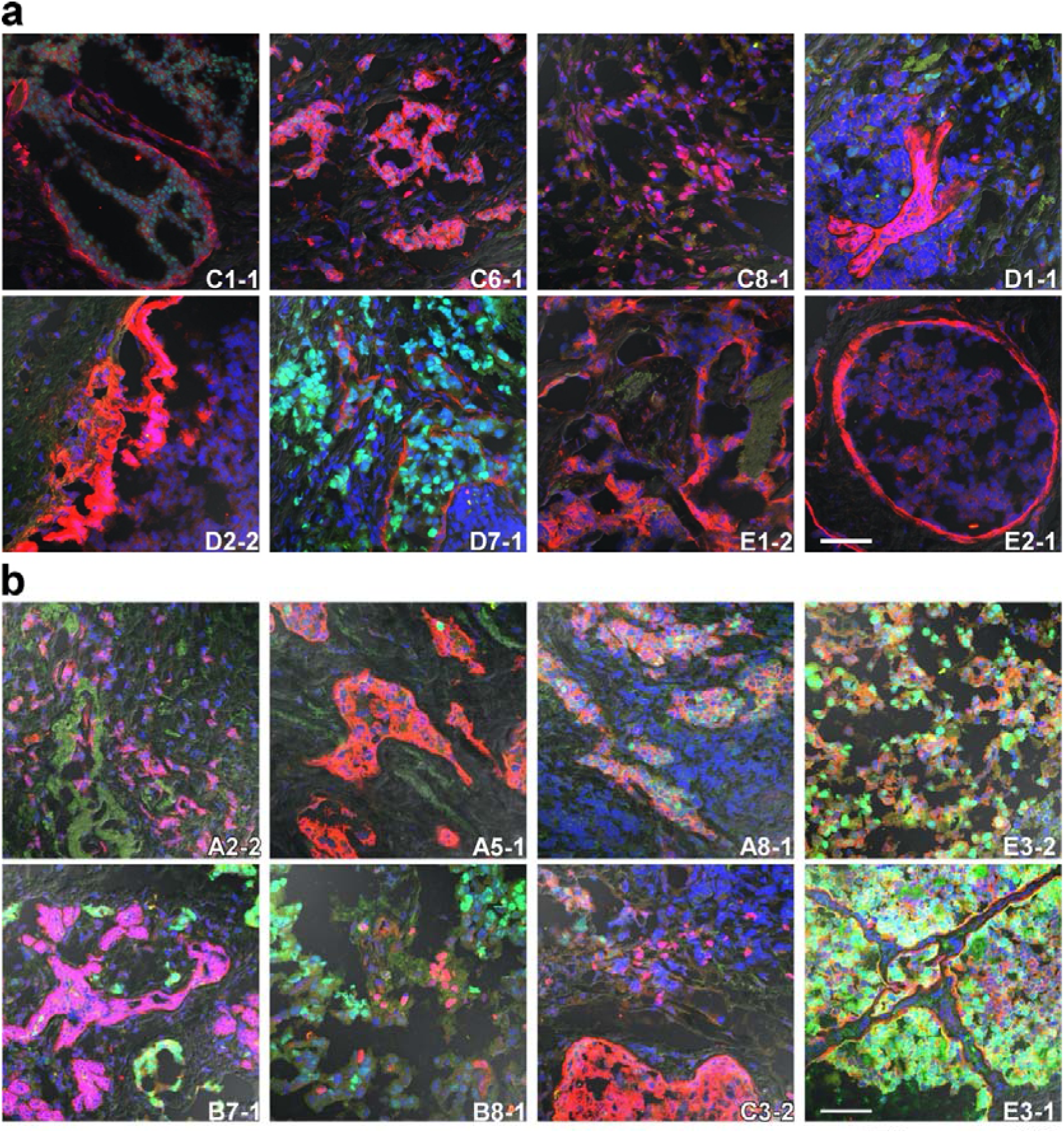
Staining patterns of breast cancer tissues with the TNBC89m clone. Representative images of tissue staining were merged; clones are indicated in red. **a**, ER, green colour. **b**, PgR, green colour. Sample IDs and observed field numbers are inset. Scale bars = 63.7 μm. Unmerged images are shown in Supplementary Figure 7.

## Discussion

We generated nanobodies for unknown markers of TNBC in a retrospective way, by immunizing alpacas with TNBC cell lines and enriching nanobody-expressing phages via TNBC cell-based biopanning. We analysed 17 nanobodies expressed on phages enriched with biopanning; 14 clones stained the cultured TNBC cells. Seven clones, TNBC68, TNBC89, TNBC143, TNBC242, TNBC610, TNBC673, and TNBC13289, were confirmed to stain malignant breast cancer tissues. The clones detected basal membrane and basal membrane-like regions that were distinct from apical cells expressing Her2; they also recognized sporadic cells surrounded by the Her2-overexpressing cells. Of note, a few clones exclusively stained dividing cultured cells and stained aggressive cells with nuclei containing a high amount of DNA in ductal cells of invasive ductal carcinoma tissue specimens. On the basis of these observations, we conclude that the clones recognize TNBC cells in cultured cell lines and Her2-negative cells in heterogeneous breast cancers.

Although no antigens have yet been determined, we suppose that the epitopes of the clones are different from each other because the staining patterns and cells of the clones are not the same. The TNBC89 and TNBC143 clones stained the broadest spectrum of cultured TNBC cell lines. TNBC242 stained cultured TNBC cell lines as well as Her2-positive SK-BR-3 cells; it stained not only Her2-positive cells but also other Her2-negative cells in breast cancer tissues. TNBC259 significantly stained cultured TNBC cell lines but not breast cancer tissues. Similarly, TNBC21 slightly stained some of the cultured TNBC cell lines but not breast cancer tissues. TNBC68 did not stain cultured TNBC cell lines well; however, it stained breast cancer tissues strongly. The TNBC13289 clone stained three basal-like TNBC cell lines but not four mesenchymal TNBC cell lines. Among the breast cancer tissue samples tested, the TNBC13289 clone stained only some cells that did not express Her2. We expect that sorting cells from breast cancer tissues using TNBC13289 could gather basal-like cells^33, 34^. It has been statistically confirmed that the prognosis of patients who are diagnosed with basal-like TNBC is worse than that of patients with other breast cancer types^29^. TNBC13289 has the potential to serve as a diagnostic antibody for surveying basal-like TNBC patients; it could also be used for CAR-T-cell therapy^35^.

## Materials and Methods

### Ethics statement

All animal experiments on alpacas were conducted in accordance with the KYODOKEN Institute for Animal Science Research and Development (Kyoto, Japan). Veterinarians performed breeding, health maintenance, and immunization studies by adhering to the published Guidelines for Proper Conduct of Animal Experiments by the Science Council of Japan. The KYODOKEN Institutional Animal Care and Use Committee approved the protocols for these studies (KYODOKEN protocol number 20200312). One alpaca (Vicugna pacos)—a 5-year-old female named Melody—was immunized, and blood samples were collected under anaesthesia.

### Immunohistochemistry

Frozen tissue sections, including breast cancer and healthy tissues, were purchased from BioChain. The embedded frozen tissue sections were fixed and permeabilized in 2% PFA and 0.05% Triton X-100 in PBS (PBST) for 20 min and further blocked with 1% goat serum in PBS (blocking buffer) at 4°C overnight. The blocked tissues were then incubated with the anti-Her2 antibody (Herceptin, mouse monoclonal, 1:300 dilution ratio) and each His-tagged nanobody at a concentration of 3 μg ml^-1^ in blocking buffer at 4°C overnight. The specimens were washed 3 times with PBST and incubated with secondary antibodies (anti-His antibody, rabbit monoclonal, 1:1000 dilution ratio) at room temperature for 1 h. After 3 washes with PBST, the slides were incubated with Alexa Fluor 488 anti-mouse and Alexa Fluor 594 anti-rabbit ternary antibodies (1:3000 dilution ratio) in PBST at room temperature for 1 h. The nuclei were labelled with DAPI.

### Image acquisition and scoring

The tissue sections were imaged using a laser scanning confocal microscope (FV1200, Olympus) with a 40x (UPLSAPO 40XSIR, 1.40 NA, Olympus) objective lens. The microscope was configured with four lasers, diode (440 nm), argon (488 nm), diode-pumped solid-state (560 nm), and helium-neon (633 nm). Images acquired with the 40× objective lens had a bit depth of 16 bits and a frame size of 1,024 ×1,024 pixels, with an image pixel size of 0.128 μm or 0.311 μm.

Fluorescence colours were detected using FV-EZ software. The resulting spectra were stored in the spectrum database of the microscope. Linear unmixing separated mixed signals pixel by pixel using proprietary software. This algorithm encompasses the entire emission spectrum of each fluorescent probe in the sample.

A blinded observer evaluated the intensity of the microscope images using a semiquantitative scoring system. The percentage of positive cells was categorized as 0 (negative staining in all fields), 1 (< 1% of cells stained), 2 (1-10% cells stained), 3 (10-50% cells stained), or 4 (>50% cells stained), and staining intensity was categorized as 0 (negative), 1 (weak), 2 (moderate), or 3 (strong). Adding the two scores together yielded a maximum score of 7.

### Antigen preparation

The immunization antigens were PFA-fixed cultured cells. Cultured breast cancer cell lines SK-BR-3, MDA-MB-231, T47D, BT-474, HS578T, BT-549, MDA-MB-436, MDA-MB-468, and HCC1937 were harvested in ice-cold PBS containing 2 mM EDTA by pipetting. The collected cells were fixed with 2% PFA at 4°C for at least 2 h. The fixed cells were washed with PBS three times and suspended in PBST (PBS with 0.01% Tween-80, Nacalai Tesque).

### Immunization and sample collection

The prepared antigen was subcutaneously injected into the alpacas five times at 2-week intervals. Blood samples were collected from the jugular vein. Peripheral blood mononuclear cells were obtained with a sucrose density gradient using Ficoll (Nacalai Tesque), washed with PBS, suspended in RNAlater solution (Thermo Fisher Scientific K.K., Tokyo, Japan), and stored in a freezer until use. Total RNA was isolated from the PBMC samples using the Direct-Zol RNA MiniPrep kit (Zymo Research, Irvine, CA).

### Library generation

Complementary DNA was synthesized from up to 1 μg of total RNA as a template with random hexamer primers and using SuperScript III reverse transcriptase (Thermo).

Coding regions of the heavy-chain variable domains were amplified with two PAGE-purified primers (CALL001: 5’-GTCCTGGCTGCTCTTCTACAAGG-3’ and CALL002: 5’-GGTACGTGCTGTTGAACTGTTCC-3’) using LA Taq polymerase (TAKARA Bio Inc., Shiga, Japan). The amplified fragments were separated on a 1.5% low-melting-temperature agarose gel (Lonza Group AG, Basel, Switzerland), and lower bands corresponding to the heavy-chain only immunoglobulin were extracted using the QIAquick Gel Extraction Kit (Qiagen K.K., Tokyo, Japan). Subsequent nested PCR was performed to amplify coding genes of VHH domains using the VHH-PstI-For and VHH-BstEII-Rev primers and subcloned into the pMES4 phagemid vector^36^. Electroporation-competent *Escherichia coli* TG1 cells (Agilent Technologies Japan, Ltd., Tokyo, Japan) were transformed with the ligated plasmids using the Gene Pulser instrument (Bio-Rad Laboratories, Inc., Hercules, CA). The library densities were monitored and maintained at >10^7^ colony-forming units per microlitre with limiting dilution. Colonies from 8 ml of cultured *E. coli* cells were harvested, pooled, and reserved in 50% glycerol stocks in a deep freezer as mother libraries.

### Plasmid construction

The whole VHH gene was synthesized into the pcDNA3.1(+) vector with the ILco2 signal peptide and the 6×His-tag. Two PstI sites on the vector were mutated with a site-direct mutagenesis kit (Agilent)—PstI-free pcDNA3.1(+) vector. Subsequently, the synthesized VHH genes were subcloned into the PstI-free pcDNA3.1(+) vector using the PstI and BstEII sites.

### Biopanning

Two rounds of biopanning were performed using PFA-fixed cultured cells. For the first round, phages dissolved in PBS containing 0.05% Tween 20, 0.5 mM NaCl, and 0.3% bovine serum albumin (BSA, Nacalai) were absorbed into PFA-fixed control cells including HEK293, HeLa, A-431, COS-7, and HepG2, for 10 min at room temperature.

Unbound phages were collected and reacted with PFA-fixed TNBC cell lines including MDA-MB-231, T47D, BT-474, HS578T, BT-549, MDA-MB-436, MDA-MB-468, HCC1937, or HCC1599, for 30 min at room temperature. The cells were washed with PBS containing 0.05% Tween 20, 0.5 mM NaCl, and 0.3% BSA 3 times; the remaining phages bound to the cells were eluted with a trypsin-ethylenediaminetetraacetic acid (EDTA) solution (Nacalai) at room temperature for 30 min with gentle agitation. The elute was neutralized with a PBS-diluted protein inhibitor cocktail (c*O*mplete, EDTA-free, protease inhibitor cocktail tablets: Roche Diagnostics GmbH, Mannheim, Germany) and used to infect TG1 cells. The infected *E. coli* was cultured and selected in LB Miller broth containing 100 μg ml^-1^ ampicillin at 37°C overnight. The selected phagemids were collected using a QIAprep Miniprep Kit (Qiagen).

### Amplicon sequencing

The VHH-coding regions within the parent (mother) libraries and the target-panned sublibraries were PCR amplified and purified using AMPure XP beads (Beckman Coulter, High Wycombe, UK). Dual-indexed libraries were prepared and sequenced on an Illumina MiSeq^37^ (Illumina, San Diego, CA). More than 100,000 pairs of each library were read (Bioengineering Lab. Co., Ltd., Kanagawa, Japan). The raw read data were trimmed to remove adapter sequence^38^ (cutadapt v1.18) and low-quality reads^39^ (Trimmomatic v0.39). The remaining paired reads were merged using fastq-join^40^ and translated to amino acid sequences^41^ (EMBOSS v6.6.0.0). The VHH sequences were cropped from start to stop codons, and then the unique amino acid sequences in each library were counted using a custom Python script combining SeqKit v0.10.1^42^ and usearch v.11^43^. The binding scores of each clone, VHH_i_, were analysed by calculating the *P* value of χ^2^ tests for changes in the existing ratios in the target sublibraries with respect to the mother libraries: for the χ^2^ statistics of VHH_i_,

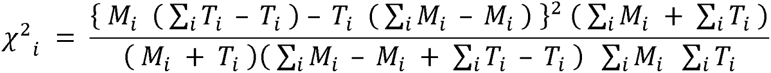

where M_i_ is the count of the VHH_i_ in the mother library, and T_i_ is the count of the VHH_i_ in the target sublibrary. The *P* value of VHH_i_ was obtained from a χ^2^ distribution with one degree of freedom. Amino acid sequences with more than 93% identity were clustered using a variant of the UCLUST algorithm^43^. The VHH clones with the most reads in each cluster that were enriched in the TNBC cell-panned sublibraries but not enriched in the negative control sublibraries were selected for further analyses.

### Antibodies

The antibodies used for immunohistochemistry (IHC) were anti-His (rabbit monoclonal EPR20547: ab213204, Abcam), anti-Her2 (mouse monoclonal anti-erbB-2: Ab00103, Absolute Antibody Ltd., Cleveland, UK), anti-ER (mouse monoclonal antibody to estrogen receptor alpha [6F11]: ab93021, Abcam), anti-PgR (mouse monoclonal anti-human progesterone receptor antibody: IHC751, GenomeMe Lab Inc., Richmond, Canada), Alexa Fluor 488 goat anti-mouse IgG (rabbit polyclonal, P0449: Dako, Glostrup, Denmark), and Alexa Fluor 594 goat anti-rabbit IgG (Dako).

### Nanobody expression

Gene sequences encoding each selected amino acid sequence were codon-optimized and synthesized (Eurofins Genomics Inc., Tokyo, Japan). Each selected amino acid sequence was expressed as a tandem dimer connected with a (GGGGS)_×4_ linker^44, 45^. The synthesized genes were subcloned into the pMES4 vector to express N-terminal PelB signal peptide-conjugated and C-terminal 6×His-tagged nanobodies^36^. BL21 (DE3) *E. coli* cells transformed with the plasmids were plated on LB agar with ampicillin and incubated at 37°C overnight. Grown colonies were picked and cultured at 37°C to reach an OD of 0.6. AU, and the cells were cultured at 37°C for 3 h with 1 mM IPTG (isopropyl-β-D-thiogalactopyranoside, Nacalai). The cultured cells were pelleted by centrifugation and stored in a freezer until use. Nanobodies were eluted from the periplasm by soaking in TES buffer (200 mM Tris, 0.125 mM EDTA, 125 mM sucrose, and pH 8.0) at 4°C for 1 h. They were further incubated with a 2× volume of 0.25× diluted TES buffer with a trace amount of benzonase nuclease (Merck) at 4°C for 45 min.

The supernatants were centrifuged (20,000×g, 4°C for 10 min), sterilized with the addition of gentamicin (Thermo), and passed through a 0.22-μm filter (Sartorius AG, Gottingen, Germany). The filtered supernatants were applied to a HisTrap HP nickel column (Cytiva) equipped on an ÄKTA pure HPLC system; the bound His-tagged nanobodies were eluted with 300 mM imidazole. The eluted fractions were collected and concentrated with a VIVAspin 3,000-molecular weight cut-off filter column (Sartorius) and applied to a Superdex75 10/300 GL gel-filtration column (Cytiva) equipped on an ÄKTA pure HPLC system. Protein purity was measured via Coomassie brilliant blue (CBB)-staining (Rapid Stain CBB Kit, Nacalai).

### Cell culture

HEK293T, HeLa, COS-7, A-431, HepG2, SK-BR-3, T47D, and HS578T breast cancer cells were grown in Dulbecco’s modified Eagle’s medium (Thermo) supplemented with 10% foetal bovine serum (FBS) and antibiotics (1% penicillin and streptomycin). The cells were cultured in a humidified incubator with 5% CO_2_ at 37°C. The breast cancer cell lines SK-BR-3 (ATCC: HTB-30), MDA-MB-231 (ATCC: HTB-26), T47D (ATCC: HTB-133), BT-474 (ATCC: HTB-20), HS578T (ATCC: HTB-126), BT-549 (ATCC: HTB-122), MDA-MB-436 (ATCC: HTB-130), MDA-MB-468 (ATCC: HTB-132), HCC1937 (ATCC: CRL-2336), and HCC1599 (ATCC: CRL-2331) were from the Japanese Collection of Research Bioresources Cell Bank (JCRB), Osaka, Japan, and American Type Culture Collection (ATCC). HEK293T, HeLa (JCRB9004), COS-7 (JCRB9127), A-431 (JCRB0004), and HepG2 (JCRB1054) cells were obtained from JCRB. MDA-MB-231, MDA-MB-436, and MDA-MB-468 cells were grown in Leibovitz’s L-15 medium supplemented with 10% FBS and antibiotics (1% penicillin and streptomycin). BT-474 cells were grown in Hybri-Care medium supplemented with 10% FBS, 1.5 g l^-1^ sodium bicarbonate, and antibiotics (1% penicillin and streptomycin). BT-549, HCC1937, and HCC1599 cells were grown in RPMI-1640 medium supplemented with 10% FBS and antibiotics (1% penicillin and streptomycin).

### Microscopy analysis

Cultured cells were seeded on a 96-well culture plate (IWAKI, AGC TECHNO GLASS CO., Ltd., Shizuoka, Japan), cultured for 24 h, and then fixed with 2% PFA at 4°C overnight. After 3 washes with PBST (0.005% Tween 20), the cells were blocked with PBST containing 2% goat serum (blocking buffer) at room temperature for 1 h. Each well was soaked with 100 μl of blocking solution containing 30 ng of purified nanobody at 4°C overnight. After 3 washes with PBST, the 1:400-diluted anti-His-tagged antibody in blocking buffer was added and reacted at room temperature for 1 h. After 3 washes, Alexa Fluor anti-rabbit IgG (594 nm emission) antibody at 1:400 dilution in blocking buffer was added to the wells. The cells were labelled at room temperature for 1 h before washing 3 times with PBST. Finally, the cell nuclei were visualized with 4’,6-diamidino-2-phenylindole (DAPI). Stained cells were imaged with a 4-ms exposure time (594 nm emission) or with a 1-ms exposure time (DAPI) using an IX71S1F-3 microscope (Olympus Corporation, Tokyo, Japan) with the cellSens Standard 1.11 application (Olympus). Each full observed field corresponding to a 165 μm × 220 μm square was photographed.

### Statistics and Reproducibility

The number of replicates is indicated in the figure legend for each experiment.

## Reporting summary

Further information on the research design is available in the Nature Research Reporting Summary linked to this article.

## Data availability

Raw data, original images, and the source data used to generate the graphs are available from the corresponding author on reasonable request.

## Materials availability

Reasonable requests for materials for research use should be directed to A.I.

## Acknowledgements

We thank all the staff of COGNANO Inc., Kaori Yurugi, Naomi Tsutsui, and Takayuki Watanabe for their efforts and Hirofumi Tsuruta, especially for the expert advice in statistics. Drs. Yoshihito Tsuji, Yasato Komatsu, and Hideya Komatsu kindly discussed this project. We are pleased to thank Nobuo Nakanishi, DVM, and the veterinarians of KYODOKEN Inc. for animal care and experiments. This study was supported by POPURI Pharmacy Co., Ltd. (Kyoto, Japan) and grants from KYOTO Industrial Support Organization 21 as subsidies (Sangakuko-no-Mori and the regional industry promotion for the next generation). We also faithfully thank Satoshi Morikawa of Yamato Scientific, Co., Ltd. (Tokyo, Japan) and Ryusuke Honjo of Green Core, Co., Ltd. (Tokyo, Japan) for their support and encouragement. In addition, Drs. Hisashi Tanaka, Farnaz Dadmanesh, and Armando Giuliano of Cedars Sinai Medical Centre (CA) gave us inspiration for the significance of TNBC.

## Author contributions

A.I. conceived the study. R.M. and A.I. performed the experiments. H.Y. performed bioinformatic analyses. R.M. drew figures and wrote the first draft of the manuscript. R.K., S.Y., and K.K. prepared and certified materials. R.M., H.Y., and A.I. analysed the data. H.Y. and A.I. reviewed and commented on the manuscript. All authors approved the reviewed manuscript.

## Additional information

Supplementary information: The online version includes the available supplementary material.

Correspondence and requests for materials should be addressed to Akihiro Imura.

## Supplementary figure legends

**Figure S1.**
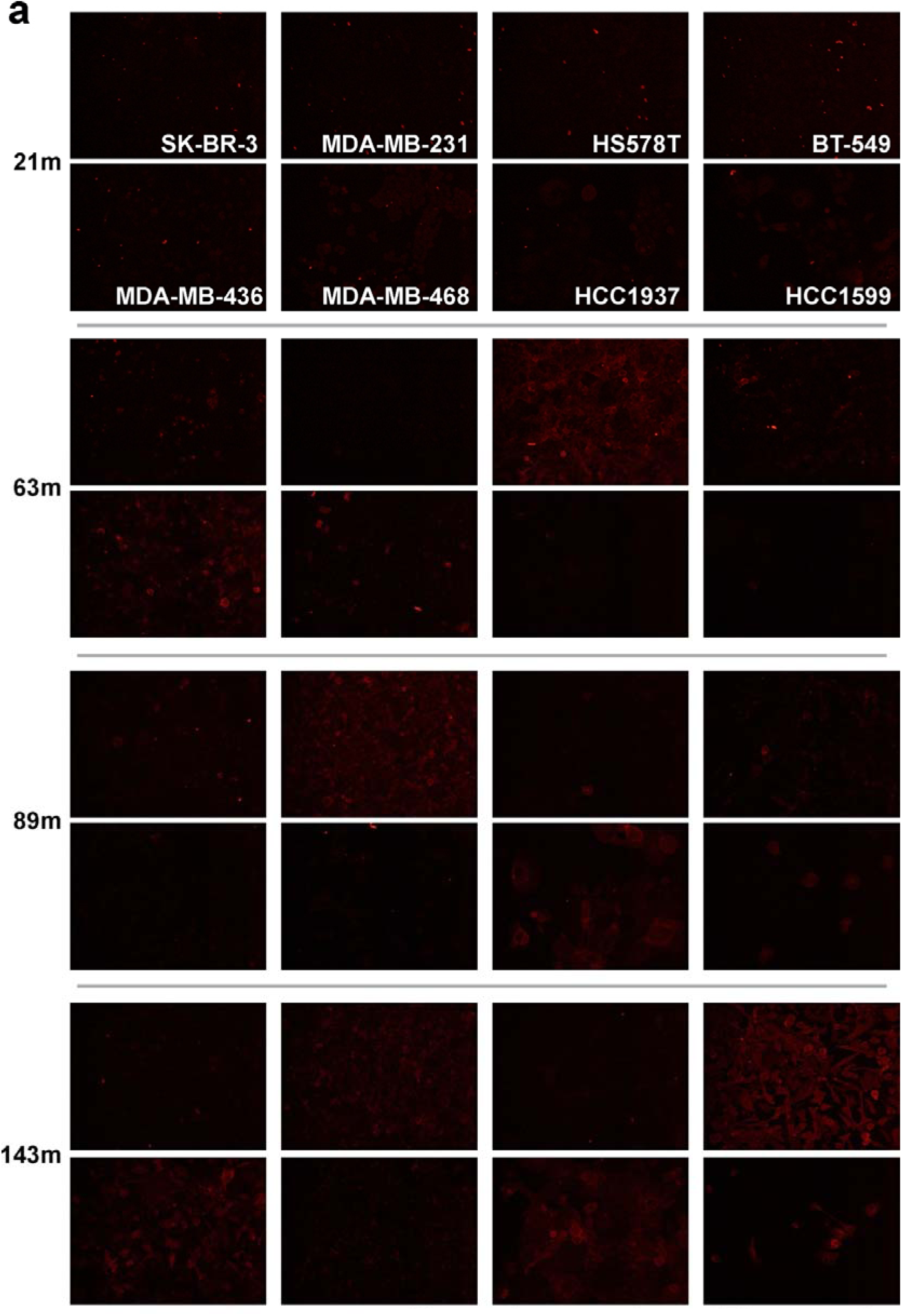
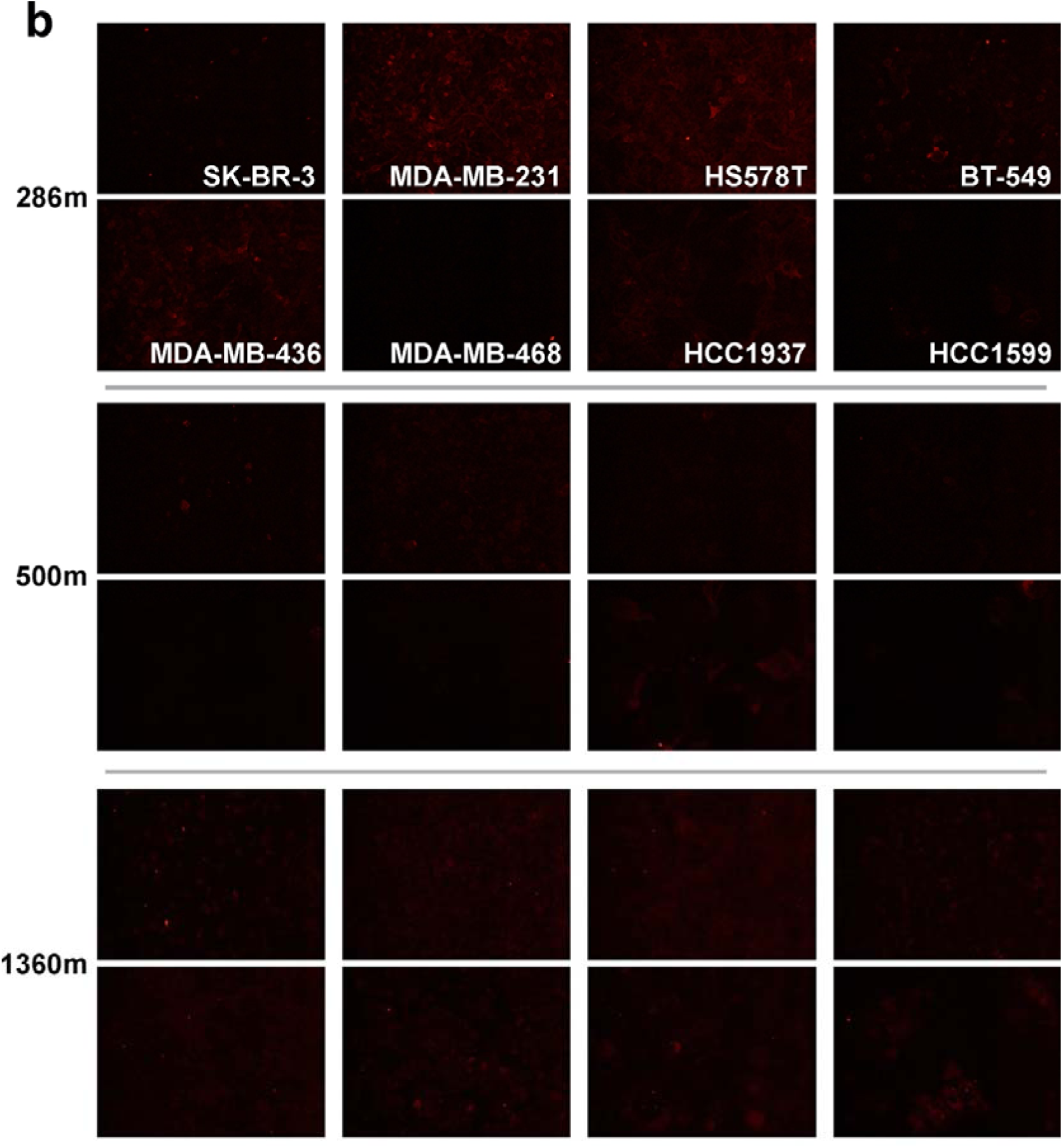
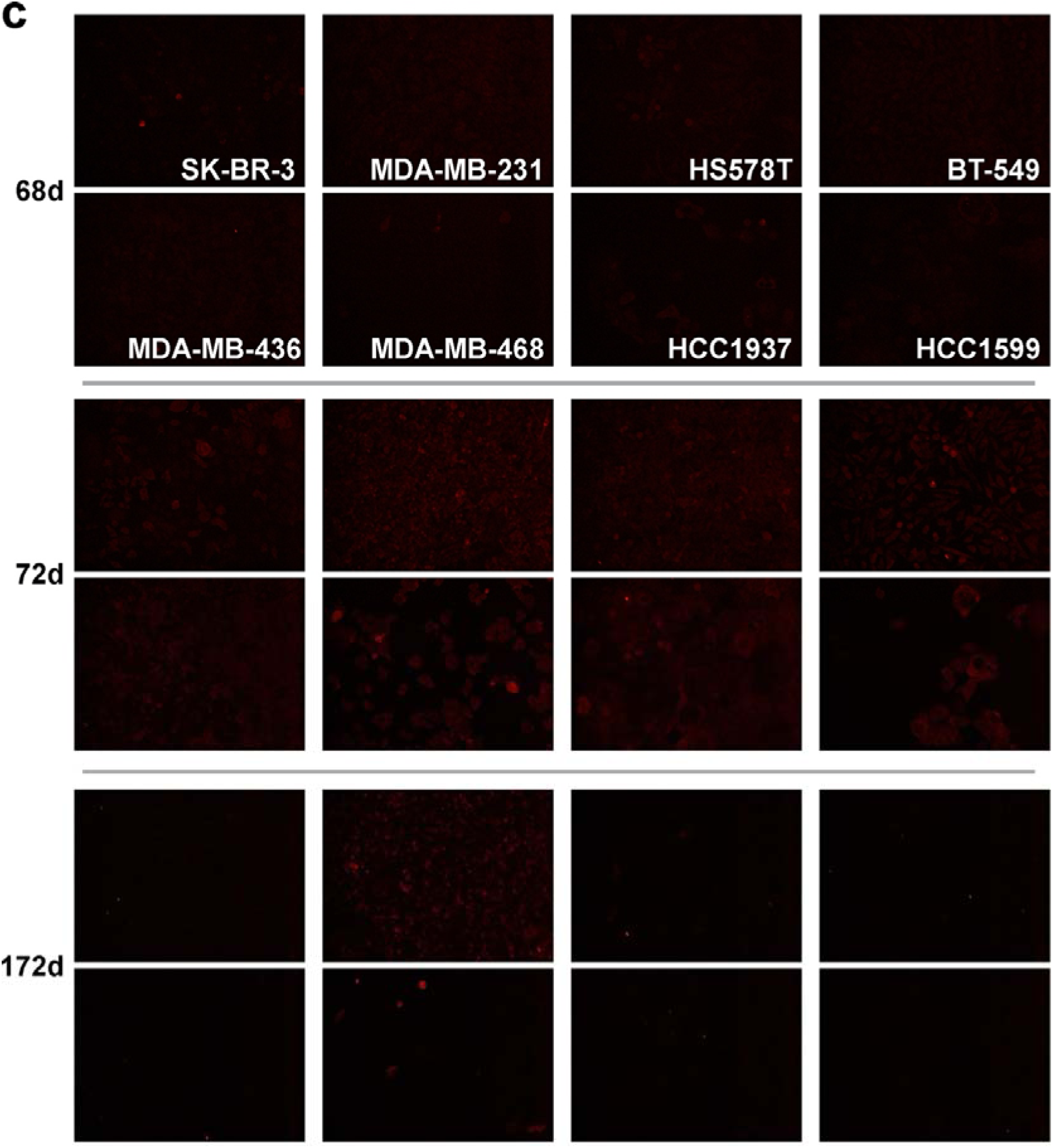
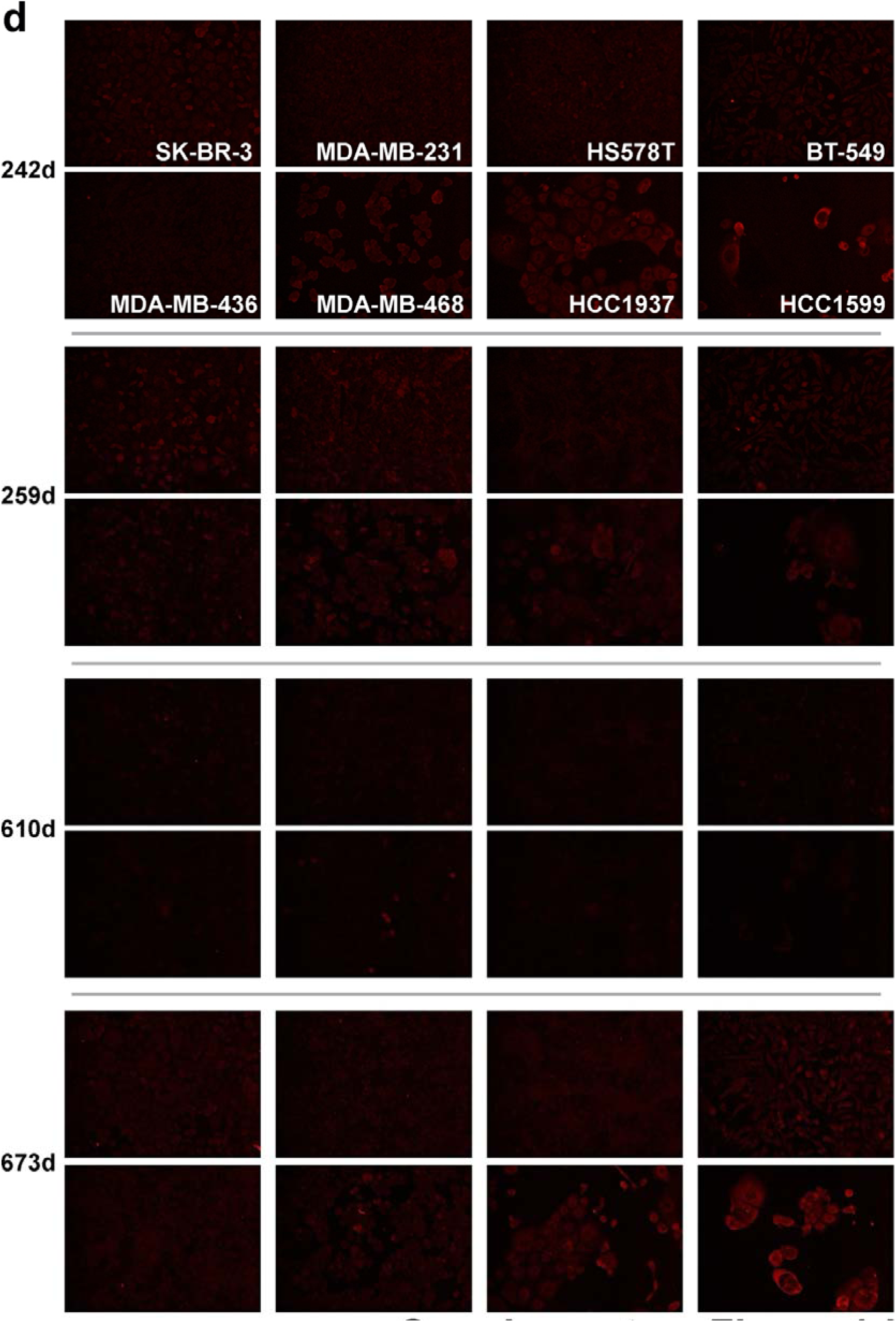
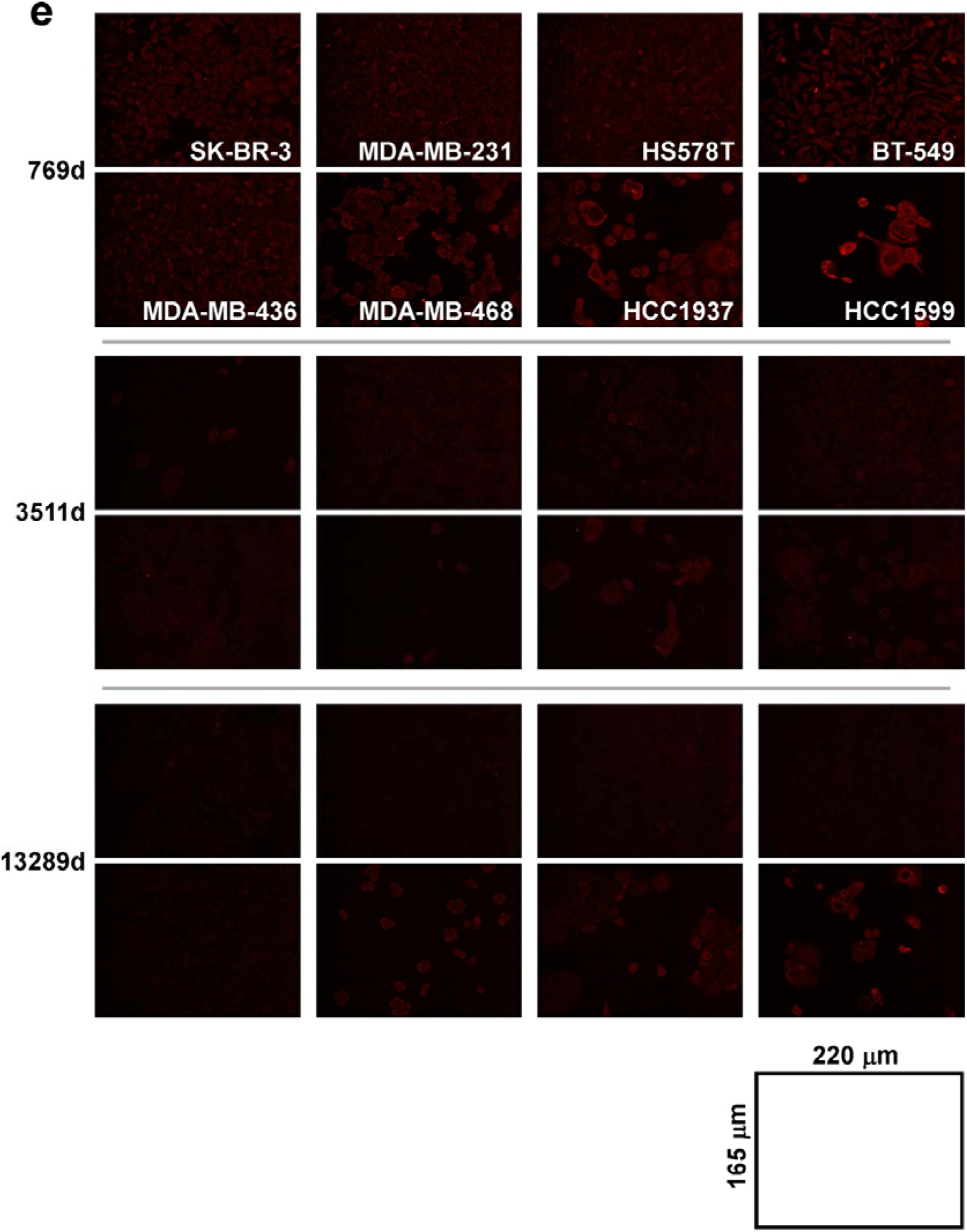
Staining of cell lines. Raw microscope images. Scales are as indicated.

**Figure S2.**
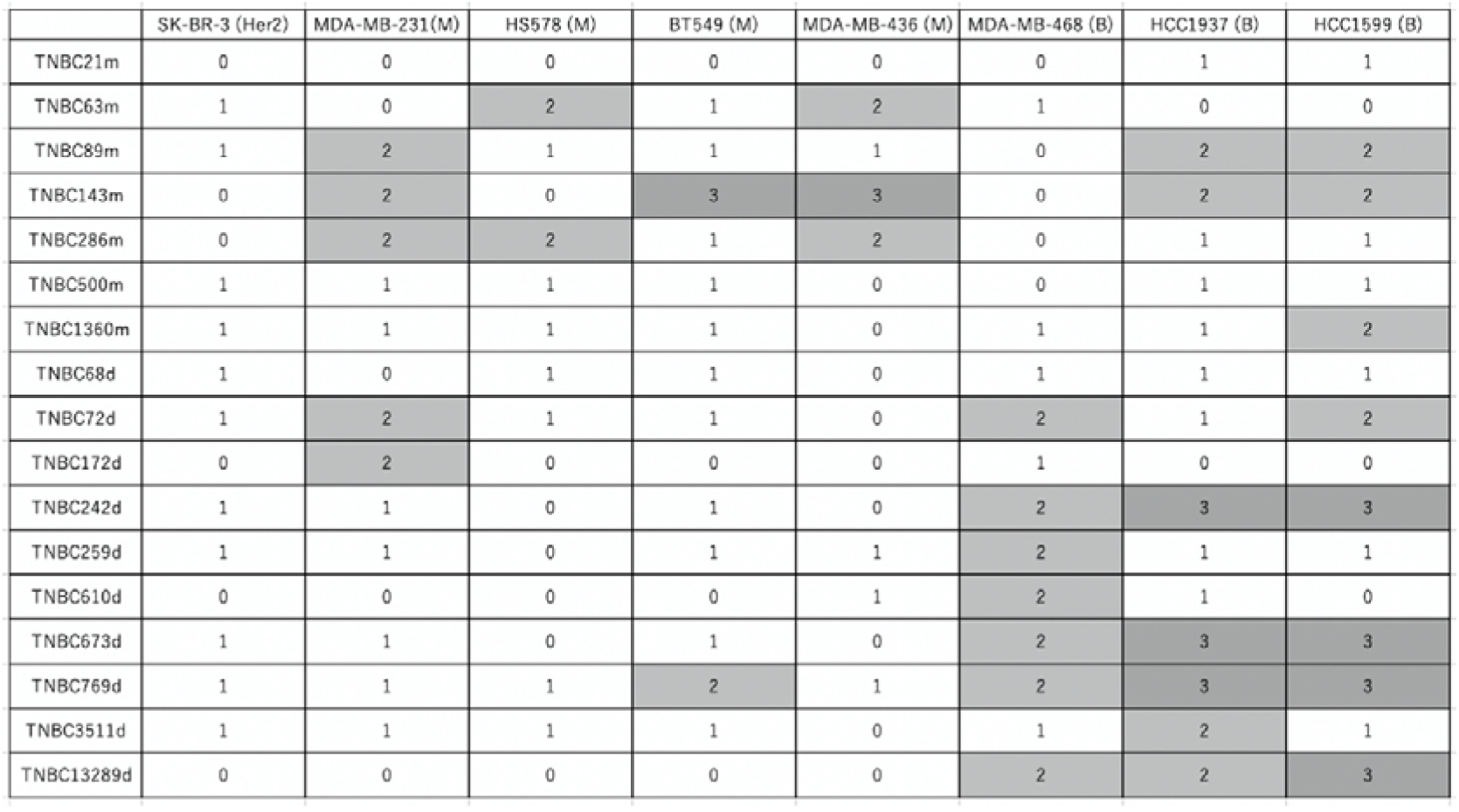
Features and staining of the breast cancer cell lines. TNBC subtypes were defined according to Lehmann et al^29^. B, basal-like; M, mesenchymal; d, dimer; m, monomer. Staining scores are shown: 3, strongly positive; 2, moderately positive; 1, weakly positive, partially (<10%) positive, or only dividing cells positive; 0, negative.

**Figure S3.**
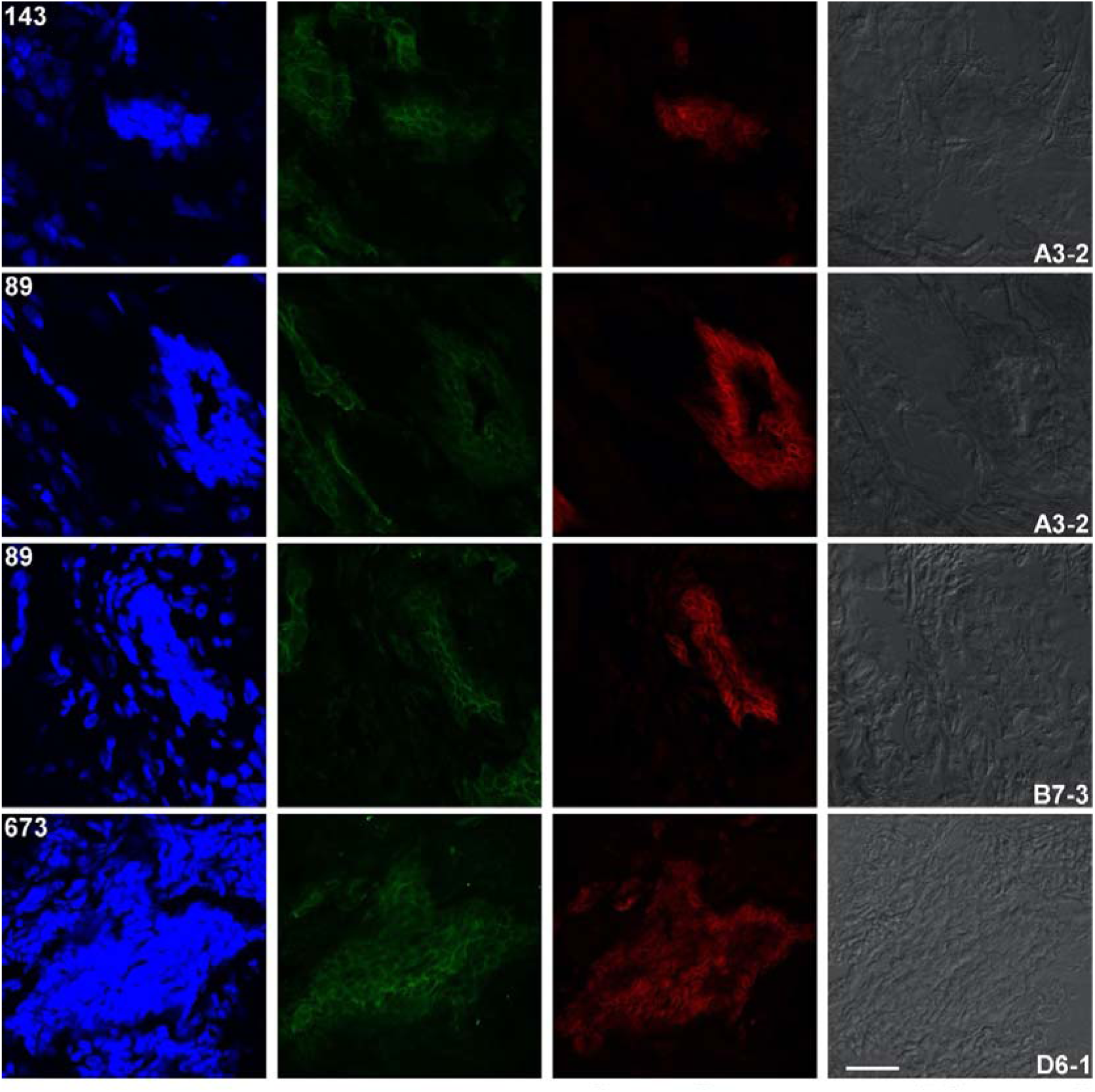
Separated image data for Figures 3, 4, and 5.

**Figure S4.**
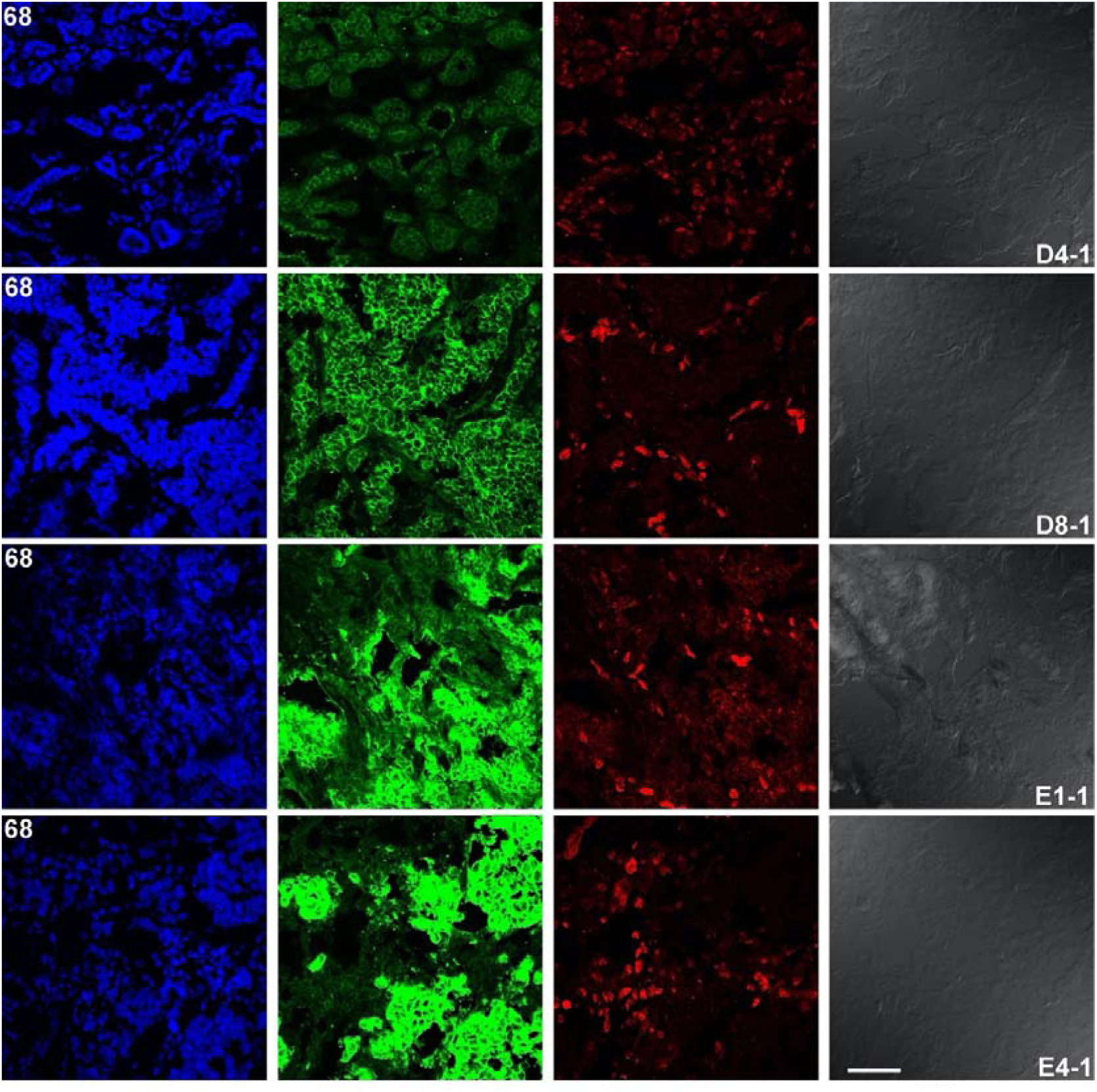
Separated image data for Figure 6.

**Figure S5.**
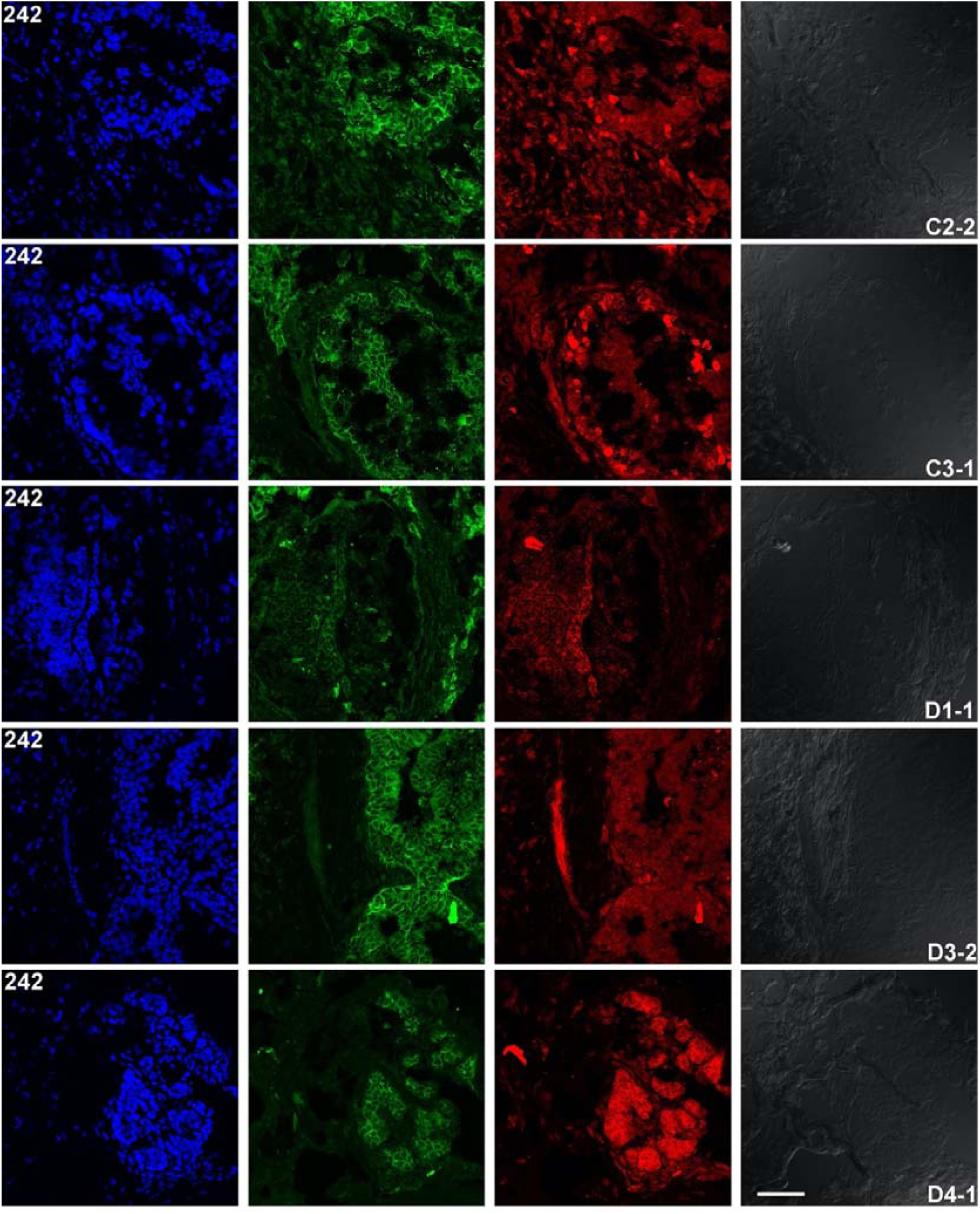
Separated image data for Figure 7.

**Figure S6.**
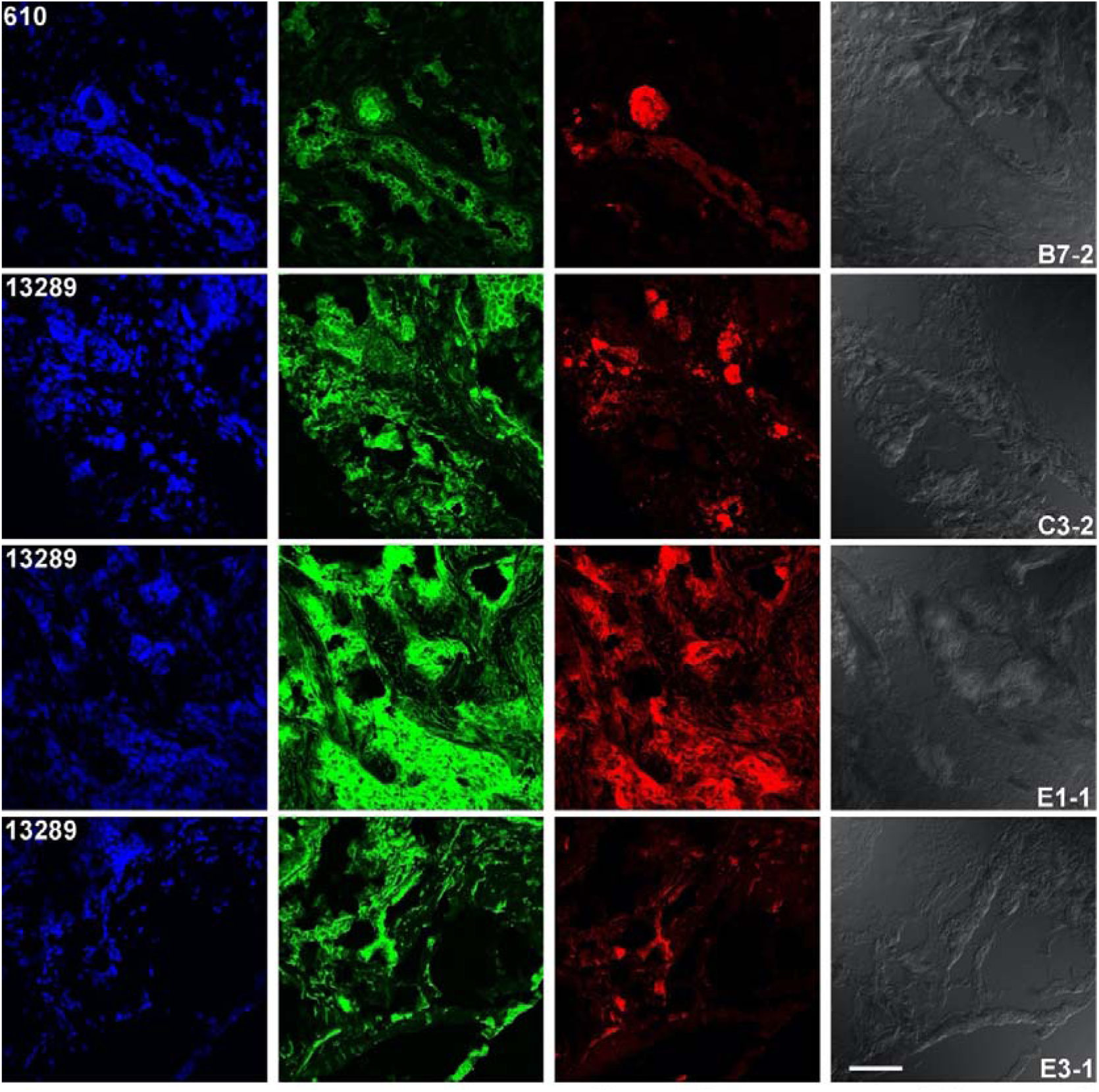
Separated image data for Figures 8 and 9.

**Figure S7.**
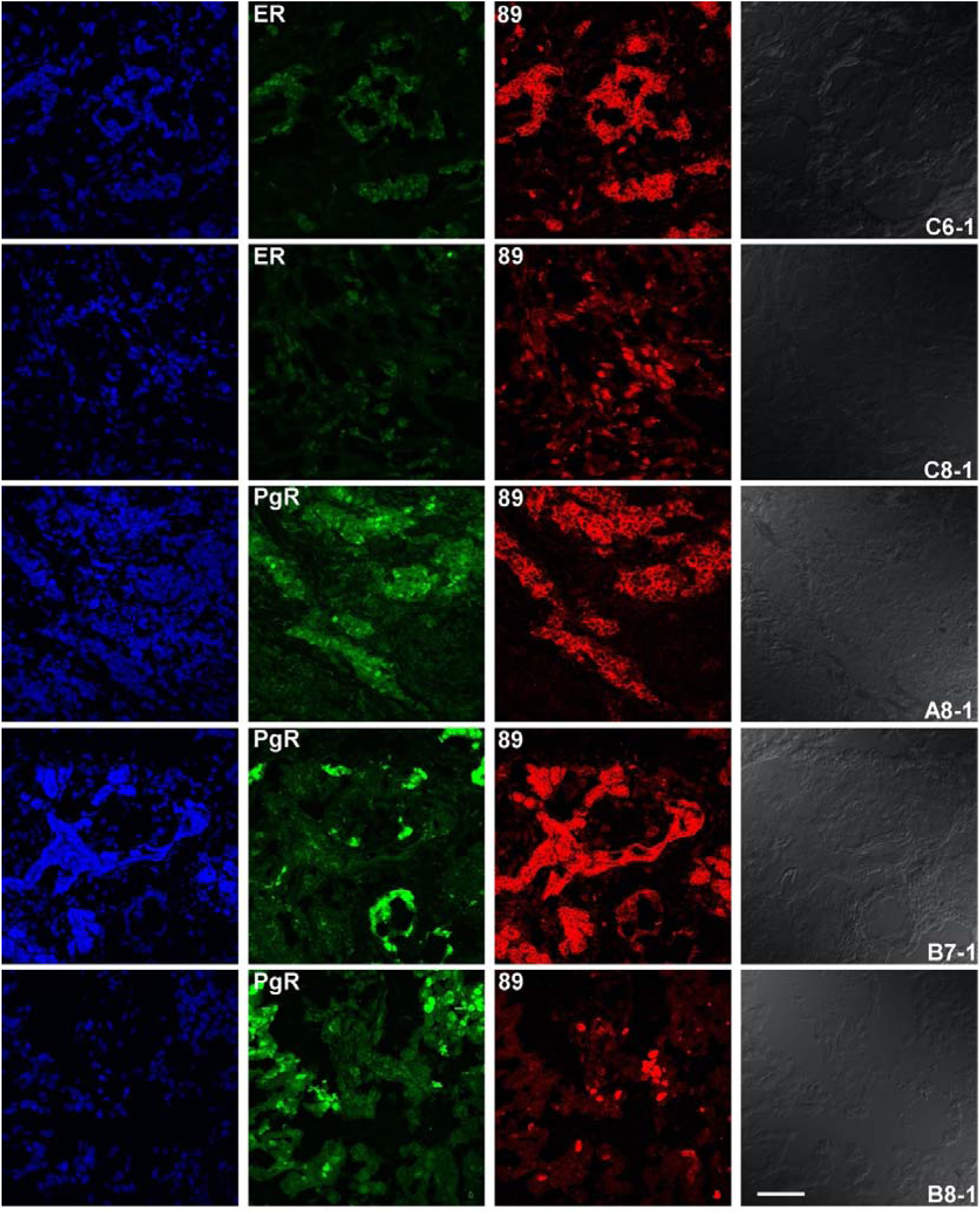
Separated image data for Figure 10.

## References

1. Siegel, R. L., Miller, K. D., Fuchs, H. E. & Jemal, A. Cancer statistics, 2022. CA Cancer J Clin 72, 7–33, doi:10.3322/caac.21708 (2022).

2. Onozato, M. L. et al. Highly Multiplexed Fluorescence in Situ Hybridization for in Situ Genomics. J Mol Diagn 21, 390–407, doi:10.1016/j.jmoldx.2019.01.010 (2019).

3. Cheng, J. et al. Clinical Validation of a Cell-Free DNA Gene Panel. J Mol Diagn 21, 632–645, doi:10.1016/j.jmoldx.2019.02.008 (2019).

4. Nielsen, T. O. et al. Immunohistochemical and clinical characterization of the basal-like subtype of invasive breast carcinoma. Clin Cancer Res 10, 5367–5374, doi:10.1158/1078-0432.CCR-04-0220 (2004).

5. Xu, J. et al. Phase II trial of veliparib and temozolomide in metastatic breast cancer patients with and without BRCA1/2 mutations. Breast Cancer Res Treat 189, 641–651, doi:10.1007/s10549-021-06292-7 (2021).

6. Suzuki, R. et al. The fragility of a structurally diverse duplication block triggers recurrent genomic amplification. Nucleic Acids Res 49, 244–256, doi:10.1093/nar/gkaa1136 (2021).

7. Harris, L. N. et al. Use of Biomarkers to Guide Decisions on Adjuvant Systemic Therapy for Women With Early-Stage Invasive Breast Cancer: American Society of Clinical Oncology Clinical Practice Guideline. J Clin Oncol 34, 1134–1150, doi:10.1200/JCO.2015.65.2289 (2016).

8. Schwartz, T., Marumoto, A. D. & Giuliano, A. E. Surgical Management of the Axilla in Breast Cancer: Evolving but Still Necessary. Ann Surg Oncol 30, 1008–1013, doi:10.1245/s10434-022-12605-x (2023).

9. Giaquinto, A. N., et al. Breast Cancer Statistics, 2022. CA Cancer J Clin 72, 524–541, doi:10.3322/caac.21754 (2022).

10. Bostrom, J. et al. Variants of the antibody herceptin that interact with HER2 and VEGF at the antigen binding site. Science 323, 1610–1614, doi:10.1126/science.1165480 (2009).

11. Vaneycken, I. et al. Preclinical screening of anti-HER2 nanobodies for molecular imaging of breast cancer. FASEB J 25, 2433–2446, doi:10.1096/fj.10-180331 (2011).

12. Roovers, R. C. et al. Efficient inhibition of EGFR signaling and of tumour growth by antagonistic anti-EFGR Nanobodies. Cancer Immunol Immunother 56, 303–317, doi:10.1007/s00262-006-0180-4 (2007).

13. Ricardo, S. et al. Breast cancer stem cell markers CD44, CD24 and ALDH1: expression distribution within intrinsic molecular subtype. J Clin Pathol 64, 937–946, doi:10.1136/jcp.2011.090456 (2011).

14. Broos, K. et al. Non-invasive assessment of murine PD-L1 levels in syngeneic tumor models by nuclear imaging with nanobody tracers. Oncotarget 8, 41932–41946, doi:10.18632/oncotarget.16708 (2017).

15. Cakir, A., Gonul, II & Uluoglu, O. A comprehensive morphological study for basal-like breast carcinomas with comparison to nonbasal-like carcinomas. Diagn Pathol 7, 145, doi:10.1186/1746-1596-7-145 (2012).

16. Jovcevska, I. et al. TRIM28 and beta-actin identified via nanobody-based reverse proteomics approach as possible human glioblastoma biomarkers. PLoS One 9, e113688, doi:10.1371/journal.pone.0113688 (2014).

17. van Brussel, A. S. et al. Hypoxia-Targeting Fluorescent Nanobodies for Optical Molecular Imaging of Pre-Invasive Breast Cancer. Mol Imaging Biol 18, 535–544, doi:10.1007/s11307-015-0909-6 (2016).

18. Duarte, J. N. et al. Generation of Immunity against Pathogens via Single-Domain Antibody-Antigen Constructs. J Immunol 197, 4838–4847, doi:10.4049/jimmunol.1600692 (2016).

19. Chatalic, K. L. et al. A Novel (1)(1)(1)In-Labeled Anti-Prostate-Specific Membrane Antigen Nanobody for Targeted SPECT/CT Imaging of Prostate Cancer. J Nucl Med 56, 1094–1099, doi:10.2967/jnumed.115.156729 (2015).

20. Rashidian, M. et al. The use of (18)F-2-fluorodeoxyglucose (FDG) to label antibody fragments for immuno-PET of pancreatic cancer. ACS Cent Sci 1, 142–147, doi:10.1021/acscentsci.5b00121 (2015).

21. Bachran, C. et al. The activity of myeloid cell-specific VHH immunotoxins is target-, epitope-, subset- and organ dependent. Sci Rep 7, 17916, doi:10.1038/s41598-017-17948-0 (2017).

22. Rashidian, M. et al. Noninvasive imaging of immune responses. Proc Natl Acad Sci U S A 112, 6146–6151, doi:10.1073/pnas.1502609112 (2015).

23. Verhaar, E. R., Woodham, A. W. & Ploegh, H. L. Nanobodies in cancer. Semin Immunol 52, 101425, doi:10.1016/j.smim.2020.101425 (2021).

24. Heukers, R. et al. Targeting hepatocyte growth factor receptor (Met) positive tumor cells using internalizing nanobody-decorated albumin nanoparticles. Biomaterials 35, 601–610, doi:10.1016/j.biomaterials.2013.10.001 (2014).

25. Behdani, M. et al. Generation and characterization of a functional Nanobody against the vascular endothelial growth factor receptor-2; angiogenesis cell receptor. Mol Immunol 50, 35–41, doi:10.1016/j.molimm.2011.11.013 (2012).

26. Tang, J. et al. Novel CD7-specific nanobody-based immunotoxins potently enhanced apoptosis of CD7-positive malignant cells. Oncotarget 7, 34070–34083, doi:10.18632/oncotarget.8710 (2016).

27. Samec, N. et al. Glioblastoma-specific anti-TUFM nanobody for in-vitro immunoimaging and cancer stem cell targeting. Oncotarget 9, 17282–17299, doi:10.18632/oncotarget.24629 (2018).

28. Maeda, R. et al. A panel of nanobodies recognizing conserved hidden clefts of all SARS-CoV-2 spike variants including Omicron. Commun Biol 5, 669, doi:10.1038/s42003-022-03630-3 (2022).

29. Lehmann, B. D. et al. Identification of human triple-negative breast cancer subtypes and preclinical models for selection of targeted therapies. J Clin Invest 121, 2750–2767, doi:10.1172/JCI45014 (2011).

30. Chavez, K. J., Garimella, S. V. & Lipkowitz, S. Triple negative breast cancer cell lines: one tool in the search for better treatment of triple negative breast cancer. Breast Dis 32, 35–48, doi:10.3233/BD-2010-0307 (2010).

31. Hurd, C. et al. Hormonal regulation of the p53 tumor suppressor protein in T47D human breast carcinoma cell line. J Biol Chem 270, 28507–28510, doi:10.1074/jbc.270.48.28507 (1995).

32. van Slooten, H. J. et al. Outgrowth of BT-474 human breast cancer cells in immune-deficient mice: a new in vivo model for hormone-dependent breast cancer. Br J Cancer 72, 22–30, doi:10.1038/bjc.1995.271 (1995).

33. Cassetta, L. et al. Human Tumor-Associated Macrophage and Monocyte Transcriptional Landscapes Reveal Cancer-Specific Reprogramming, Biomarkers, and Therapeutic Targets. Cancer Cell 35, 588–602 e510, doi:10.1016/j.ccell.2019.02.009 (2019).

34. Tuit, S. et al. Transcriptional Signature Derived from Murine Tumor-Associated Macrophages Correlates with Poor Outcome in Breast Cancer Patients. Cell Rep 29, 1221–1235 e1225, doi:10.1016/j.celrep.2019.09.067 (2019).

35. Han, S. et al. Recent clinical trials utilizing chimeric antigen receptor T cells therapies against solid tumors. Cancer Lett 390, 188–200, doi:10.1016/j.canlet.2016.12.037 (2017).

36. Pardon, E. et al. A general protocol for the generation of Nanobodies for structural biology. Nat Protoc 9, 674–693, doi:10.1038/nprot.2014.039 (2014).

37. Zomnir, M. G. et al. Artificial Intelligence Approach for Variant Reporting. JCO Clin Cancer Inform 2, doi:10.1200/CCI.16.00079 (2018).

38. Martin, M. Cutadapt removes adapter sequences from high-throughput sequencing reads. 2011 17, 3, doi:10.14806/ej.17.1.200 (2011).

39. Bolger, A. M., Lohse, M. & Usadel, B. Trimmomatic: a flexible trimmer for Illumina sequence data. Bioinformatics 30, 2114–2120, doi:10.1093/bioinformatics/btu170 (2014).

40. Aronesty, E. Comparison of Sequencing Utility Programs. The Open Bioinformatics Journal 7, 1–8, doi:10.2174/1875036201307010001 (2013).

41. Rice, P., Longden, I. & Bleasby, A. EMBOSS: the European Molecular Biology Open Software Suite. Trends in genetics : TIG 16, 276–277, doi:10.1016/s0168-9525(00)02024-2 (2000).

42. Shen, W., Le, S., Li, Y. & Hu, F. SeqKit: A Cross-Platform and Ultrafast Toolkit for FASTA/Q File Manipulation. PLOS ONE 11, e0163962, doi:10.1371/journal.pone.0163962 (2016).

43. Edgar, R. C. Search and clustering orders of magnitude faster than BLAST. Bioinformatics 26, 2460–2461, doi:10.1093/bioinformatics/btq461 (2010).

44. Nagata, K. et al. Intratracheal trimerized nanobody cocktail administration suppresses weight loss and prolongs survival of SARS-CoV-2 infected mice. Commun Med (Lond) 2, 152, doi:10.1038/s43856-022-00213-5 (2022).

45. Yamaguchi, K. et al. Structural insights into the rational design of a nanobody that binds with high affinity to the SARS-CoV-2 spike variant. J Biochem 173, 115–127, doi:10.1093/jb/mvac096 (2023).

